# A tissue centric atlas of cell type transcriptome enrichment signatures

**DOI:** 10.1101/2023.01.10.520698

**Authors:** P Dusart, S Öling, E Struck, M Norreen-Thorsen, M Zwahlen, K von Feilitzen, P Oksvold, M Bosic, MJ Iglesias, T Renne, J Odeberg, F Pontén, C Lindskog, M Uhlén, LM Butler

## Abstract

Genes with cell type specific expression typically encode for proteins that have cell type specific functions. Single cell RNAseq (scRNAseq) has facilitated the identification of such genes, but various challenges limit the analysis of certain cell types and lowly expressed genes. Here, we performed an integrative network analysis of over 6000 bulk RNAseq datasets from 15 human organs, to generate a tissue-by-tissue cell type enrichment prediction atlas for all protein coding genes. We profile all the major constituent cell types, including several that are fragile or difficult to process and thus absent from existing scRNAseq-based atlases. The stability and read depth of bulk RNAseq data, and the high number of biological replicates analysed, allowed us to identify lowly expressed cell type enriched genes that are difficult to classify using existing methods. We identify co-enriched gene panels shared by pancreatic alpha and beta cells, chart temporal changes in cell enrichment signatures during spermatogenesis, and reveal that cells in the hair root are a major source of skin enriched genes. In a cross-tissue analysis, we identify shared gene enrichment signatures between highly metabolic and motile cell types, and core identity profiles of cell types found in across tissue types. Our study provides the only cell type gene enrichment atlas generated independently of scRNAseq, representing a new addition to our existing toolbox of resources for the understanding of gene expression across human tissues.

## INTRODUCTION

Cell type can be categorised by function, origin, location, morphology and, more recently, global transcriptome. Transcriptional profiles depend on both intrinsic cell characteristics and transient states, but selective expression of genes typically required for cell type specialised functions currently underlie our definition of cell type. Large-scale projects, such as the Human Cell Atlas (www.humancellatlas.org) (Regev et al., 2017) and the Human Protein Atlas (www.proteinatlas.org/) (Karlsson et al., 2021; Uhlen et al., 2015) contain single-cell RNA sequencing (scRNA-seq) data from thousands of cells, which can be used to further understand human health and disease, through, for example, targeted biomarker discovery (Iglesias et al., 2021), or elucidation of disease associated gene expression (Cano-Gamez and Trynka, 2020; Wang, 2021).

However, scRNA-seq has limitations; cell processing can cause artefactual modification of gene expression, through induction of the stress response (Denisenko et al., 2020; van den Brink et al., 2017) or as a consequence of removal from the microenvironment (Massoni-Badosa et al., 2020). Some cell types are sensitive to extraction protocols, e.g., kidney podocytes (Denisenko et al., 2020), whilst others require extensive, damaging proteolytic digestion to isolate e.g., adipocytes (Rondini and Granneman, 2020; Viswanadha and Londos, 2006); such cell types are absent from the major databases (Karlsson et al., 2021; Quake, 2021; Tabula Muris et al., 2018). Single nuclei sequencing is an alternative tool for analysing such cell types (Habib et al., 2017), but resultant expression profiles are incomplete (Thrupp et al., 2020). Compared to bulk RNA-seq, where all cell types in a tissue are sequenced without prior separation, scRNAseq produces less stable and more variable data, with a high number of zero values, particularly for lowly expressed genes (Haque et al., 2017; Hicks et al., 2018; Kolodziejczyk et al., 2015; Zheng et al., 2021), requiring computational imputation for interpretation (Chen et al., 2019; Hou et al., 2020), with methods remaining controversial (Jiang et al., 2022). Typically, tissues from a limited number of donors are analysed, resulting in underestimation of biological variance of gene expression and potential false discoveries when analysing differential expression between cell types or conditions (Denninger et al., 2022; Squair et al., 2021; Trostle et al., 2022). Differentially expressed genes identified using scRNAseq typically have higher expression and smaller fold changes than those identified with bulk RNAseq (Denninger et al., 2022).

We previously developed and validated an integrative correlation analysis method to identify cell type-enriched transcriptome profiles from unfractionated tissue RNAseq (Butler et al., 2016; Dusart et al., 2019; Norreen-Thorsen et al., 2022). Our method circumvents some limitations of scRNAseq; hundreds of samples are analysed concurrently to reduce the influence of biological variation and batch effects, cell types that are technically challenging to process can be analysed, and lowly expressed cell enriched transcripts classified (Norreen-Thorsen et al., 2022). Here, we analysed over 6000 bulk RNAseq datasets from Genotype-Tissue Expression (GTEx) to generate a genome-wide, tissue-by-tissue cell type enrichment prediction atlas for all protein coding transcripts in 15 different human tissues. We provide gene enrichment signatures for all major constituent cell types, including those that are fragile or difficult to process, such as podocytes in the kidney and adipocytes in the breast, as well as for minority cell types, such as those in the hair follicles of the skin. We identify co-enriched genes shared by related cell types, such as pancreatic alpha and beta cells, and chart temporal changes in gene enrichment during spermatogenesis. In a cross-tissue analysis, we identify common gene enrichment signatures, e.g., between respiratory ciliated cells and spermatids, endocrine cells in the pancreas, colon, thyroid, and stomach, and between cell types found in all or most tissues, such as endothelial and immune cell types. All data is available on the Human Protein Atlas (HPA) (www.proteinatlas.org/humanproteome/tissue+cell+type).

## RESULTS

### Cell type reference transcripts correlate across unfractionated tissue RNAseq data

Bulk RNAseq datasets for 15 human tissue types were retrieved from Genotype-Tissue Expression (GTEx) V8 (www.gtexportal.org) (Consortium, 2015) (Figure 1A). To identify cell type-enriched transcript profiles, we performed an integrative correlation analysis on each dataset, using our previously published method (Butler et al., 2016; Dusart et al., 2019; Norreen-Thorsen et al., 2022).

**Figure 1.**
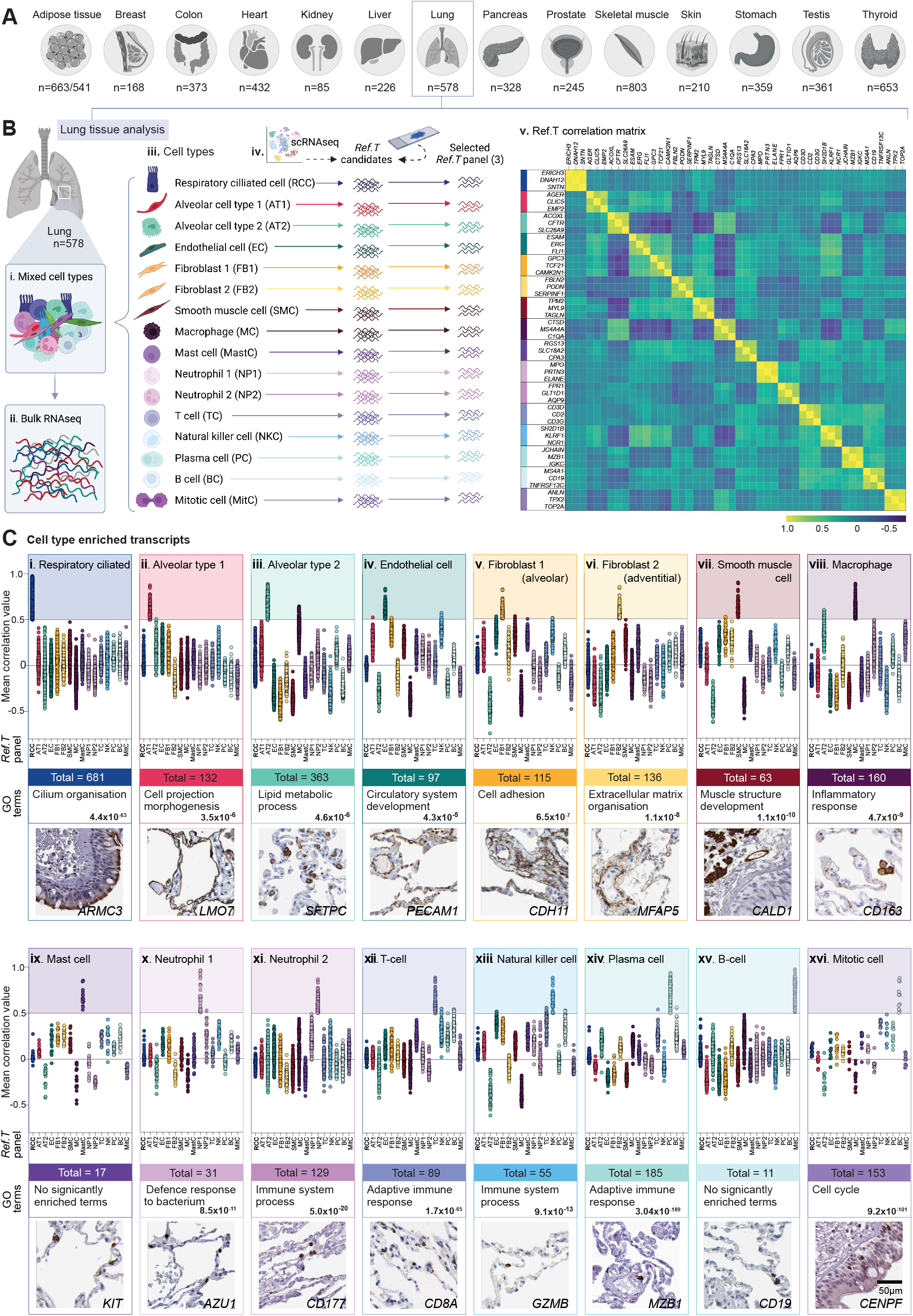
Integrative co-expression analysis of unfractionated human lung tissue RNAseq can resolve constituent cell type enriched genes. (**A**) Bulk RNAseq datasets were retrieved from GTEx V8 and analysed by tissue type (n=sample number). (**B**) Analysis concept, using lung as an illustrative example: (i) each sample (n=578) contained mixed cell types, contributing (ii) differing proportions of mRNA to each sequenced dataset. To profile cell type-enriched transcriptomes (iii) constituent cell types for each tissue were identified and for each (iv) candidate reference transcripts (*Ref.T*.) for ’virtual tagging’ were shortlisted, primarily based on predicted cell specificity from existing literature and/or in house protein profiling. (v) Matrix of correlation coefficient values between selected *Ref.T*. across the sample set. (**C**) Mean correlation coefficients between genes above designated thresholds for classification as cell-type enriched in: (i) respiratory ciliated [RCC], (ii) alveolar type I [AT1], (iii) alveolar type II [AT2], (iv) endothelial [EC], (v) alveolar fibroblasts [FB1], (vi) adventitial fibroblasts [FB2], (vii) smooth muscle cell [SMC], (viii) macrophage [MC], (ix) mast cell [MastC], (x-xi) neutrophil [NP1 and NP2], (xii) T-cell [TC], (xiii) natural killer cell [NK], (xiv) plasma cell [PC], (xv) B-cell [BC], or (xvi) mitotic cell [MitC], and all *Ref.T*. panels. Total number, most significant gene ontology (GO) terms and illustrative protein profiling in human lung tissue are provided for each cell type. See also Table S1, Figure S1 and S2.

As the tissue is unfractionated prior to sequencing, constituent cell types are present in different proportions in each sample (Figure 1 B.i [lung as an illustrative example]). Thus, each cell contributes mRNAs subsequently measured by RNAseq (Figure 1 B.ii), which can be: predominantly expressed in that cell type (cell type enriched), selectively expressed in two cell types (co-enriched), or expressed in several, or all, cell types within the tissue. For the main constituent cell types in each tissue (Figure 1 B.iii) marker ’reference transcripts’ [*Ref.T*.] were shortlisted (n=10-30), including: (**i**) those identified through in house tissue protein profiling (Uhlen et al., 2015) (**ii**) established markers identified in older ’none-omics’ studies, (**iii**) those identified by scRNAseq of mouse (Tabula Muris et al., 2018) or human (Han et al., 2020) tissue, and (**iv**) markers from databases containing multiple studies e.g., <u>Cell Marker</u> (Zhang et al., 2019), <u>PanglaoDB</u> (Franzen et al., 2019) (Figure 1 B.iv). Spearman correlation coefficients were generated between all shortlisted candidate *Ref.T*. across each sample set, and three were selected to represent each cell type (for lung see Figure 1 B.v), based on the following criteria: (**i**) a high correlation between *Ref.T*. <u>within</u> each cell type panel (FDR <0.00001), consistent with cell type co-expression, (**ii**) a low correlation coefficient between *Ref.T*. in <u>different</u> cell type panels, consistent with high *specificity* of each panel (Figure 1 B.v) and (**iii**) a normal expression distribution of *Ref.T*. across samples. For all cell types, corresponding *Ref.T* and intra/inter *Ref.T* panel correlation coefficients in each tissue see Table S1, Tab 1, Table A-O.

### Reference transcripts analysis can identify cell-type enriched gene signatures

For each tissue type analysed, the proportion of constituent cell types between samples vary, due to sampling and genetic factors (Consortium, 2020; Wang et al., 2019), but ratios between constitutively expressed cell-specific genes remain relatively constant. Thus, high correlation of a given transcript with all *Ref.T*. in any one panel is consistent with selective expression in the corresponding cell type (Norreen-Thorsen et al., 2022). For all tissues, we generated correlation coefficients between each *Ref.T*. and all other sequenced transcripts (’test-transcripts’) and produced a list of provisional cell type-enriched transcripts, based on the following criteria: (**i**) the test-transcript had a mean correlation with a given *Ref.T*. panel ≥0.50 (FDR <0.0001), which was (**ii**) higher than the mean correlation with *any other Ref.T*. panel. Resultant transcripts for each cell type were generally well separated from all others e.g., for lung: respiratory ciliated cells (RCC; Figure S1 A.i) and alveolar cell type 1 (AT1; Figure S1 Bi). However, in some cases, test-transcripts correlated well with more than one *Ref.T*. panel; panels typically representing closely related cell types, e.g., natural killer and T-cells (NK and TC; Figure S1 C.i), or those with functional commonalities, e.g., macrophages and alveolar type 2 (AT2) cells (Debbabi et al., 2005) (MC and AT2; Figure S1 D.i). To more carefully analyse the relationship between transcripts, the following was calculated for each to compare cell type lists: (**i**) the ‘*differential correlation score*’, defined as the difference between the mean correlation of the test-transcript with the two sets of *Ref.T*., e.g., respiratory ciliated cell (RCC) type panel [*ERICH3, DNAH12, SNTN*] and smooth muscle cell (SMC) panel [*TPM2, MYL9, TAGLN*] (Figure S1 A.ii) and (**ii**) the ‘*enrichment ranking*’, based on the mean correlation value of the test-transcript with the *Ref.T*. panel (rank 1 = highest corr.). Transcripts that most highly correlated with the RCC *Ref.T*. panel separated well, from even the next closest cell type, SMC (Figure S1 A.ii), as did those most highly correlating with the alveolar cell type 1 (AT1) *Ref.T*. panel (Figure S1 B.ii). A panel of transcripts that most highly correlated with *Ref.T*. representing NK (Figure S1 C.ii, right side) or MC (Figure S1 D.ii, right side) had a low differential correlation score with *Ref.T*. for TC or AT2, respectively (Figure S1 C.ii and D.ii, left side), consistent with co-enrichment in both cell types, as we previously demonstrated (Norreen-Thorsen et al., 2022). scRNAseq data from human lung (Tabula Sapiens et al., 2022) was used to verify expression profiles of selected transcripts with predicted enrichment in one (Figure S1 A-D.iii and v) or both cell types (Figure S1 A-D.iv). For classification as single cell-type enriched, any transcript with a differential correlation score <0.15 vs. any *Ref.T*. panel representing a different cell type was excluded, on the basis of predicted co-enrichment (e.g., Figure S1 A-D.ii, grey shaded area). Application of these criteria across tissues generally resulted in intra-tissue cell-enriched gene panels that were well separated from each other (example for lung; Figure 1 C.i-xvi). For some cell types, these default thresholds were decreased when overlap with other *Ref.T*. panels was absent e.g., for erythroid cells in the liver (Figure S1 E.i and ii) or increased when overlap remained (details provided in Table S1, Tab 3). Gene ontology (GO) analysis (Gene Ontology, 2021), performed to identify over-represented classes and pathways among genes identified as cell type enriched produced resultant terms consistent with expected cell type functions, e.g. for lung respiratory ciliated cells, significant terms included ’*cilium organisation*’ (FDR 4.4 ×10^−63^) (Figure 1 C.i), and for plasma cells ’*adaptive immune response*’ (FDR 3.0 ×10^−189^) (Figure 1C.xiv). Tissue profiling for selected proteins encoded by predicted cell type enriched genes had expression consistent with our classifications (Figure 1 C.i-xvi).

### Weighted network correlation analysis supports cell type enrichment predictions

As our analysis method is based on manually selected *Ref.T*., cell type classification is subject to an input bias. However, we previously showed that unbiased weighted network correlation analysis (WGCNA) (Langfelder and Horvath, 2008), where correlation coefficients between all transcripts are calculated and subsequently clustered into related groups (based on expression similarity), supports *Ref.T*. based analysis cell type enrichment predictions (Dusart et al., 2019; Norreen-Thorsen et al., 2022). Here, we performed WGNCA of lung and liver samples (Figure S2). Both *Ref.T* (Figure S2 A-B.i) and predicted cell-type enriched gene panels (Figure S2 A-B.ii-ix) clustered into the same, or closely related WGCNA groups when the differential correlation for exclusion was set at >0.15 (as described above) (Figure S2 A-B.v). When the differential correlation was increased in increments of 0.05 (Figure S2 A-B.vi-ix) the number of predicted cell type enriched genes outside the predominant WGCNA clusters decreased (see red dashed box), consistent with higher enrichment specificity. Gene enrichment could thus be categorised into *very high, high* or *moderate*, corresponding to a differential score vs. other profiled cell types within the tissue of >0.35, >0.25 or >0.15, respectively (see Table S1, Tab 3 for total number in each category for all cell types/tissues).

### Specialised cell types have the highest number of enriched genes within tissues

The total number of genes with predicted cell type enrichment (*very high, high* or *moderate*) within each tissue ranged from 7041 (testis) to 829 (pancreas) (Figure 2 A) (Table S1, Tab 3). The number of cell types analysed in each tissue type ranged from 7-18; with the lowest number profiled in skeletal muscle and subcutaneous adipose tissue (n=7 and 8, respectively) and the highest in skin and lung (n=18 and 14, respectively) (Table S1, Tab 1).Tissue specialised cell types had the highest number of enriched genes, such as cardiomyocytes in the heart (number/total enriched in all cell types in that tissue: 916/1902 [48%]) (Figure 2 B.v), proximal tubular cells in the kidney (657/1778 [37%]) (Figure 2 B.vii), hepatocytes in the liver (1264/2393 [53%]) (Figure 2 B.xi), keratinocytes in the skin (945/2460 [38.4%]) (Figure 2 B.xiii), gastric mucosal cells in the stomach (379/1361 [28%]) (Figure 2 B.xiv) and respiratory ciliated cells in the lung (681/2419 [28%]) (Figure 2 B.xv).

**Figure 2.**
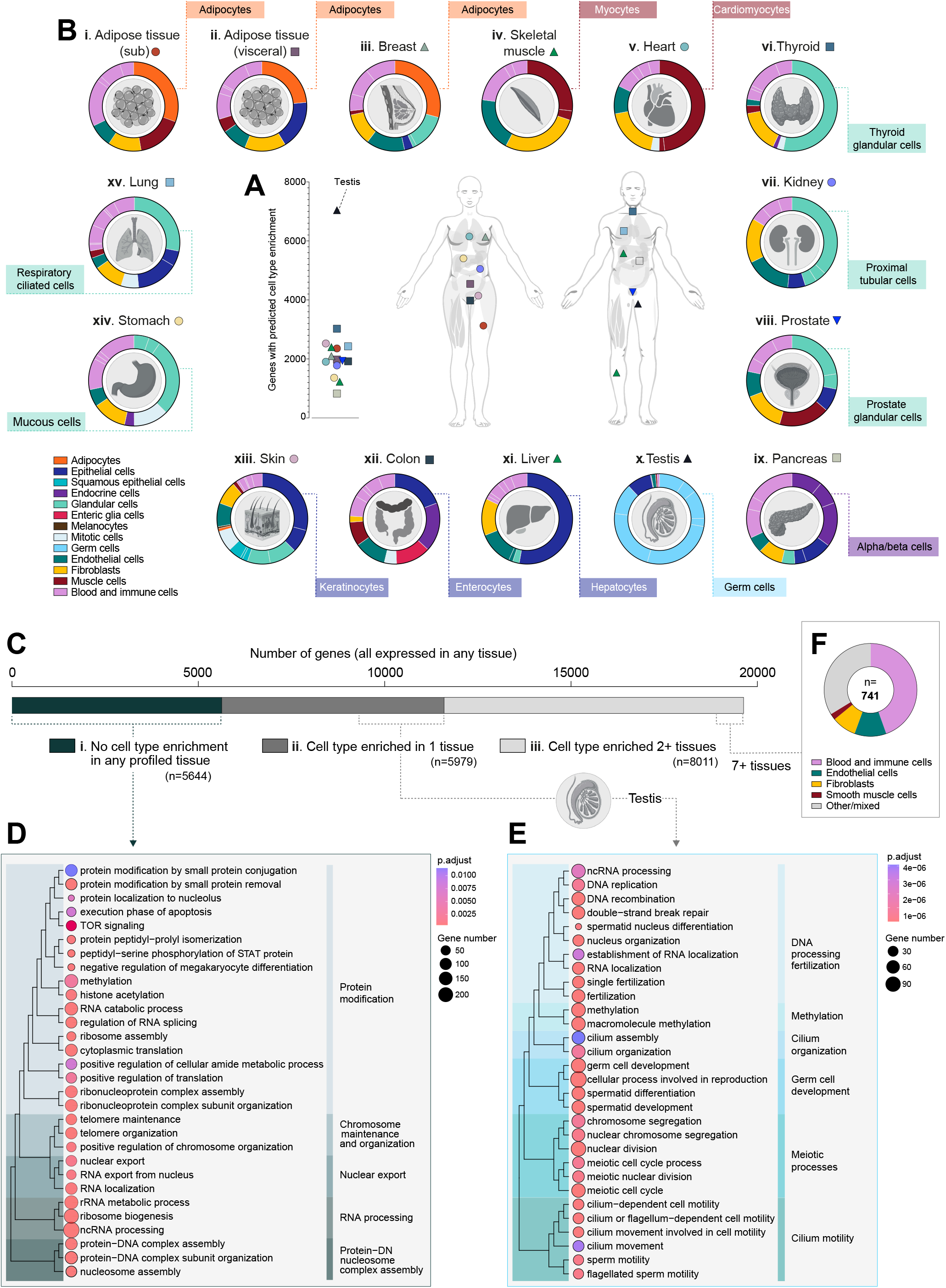
Overview of cell type enriched gene profiles across tissue types. Bulk RNAseq datasets were retrieved from GTEx V8 and cell type enriched transcriptome predictions made using integrative correlation analysis. (**A**) Number of genes with predicted cell type enrichment in each analysed tissue type. **(B)** Circular plots showing broad classification of genes predicted to be cell type enriched in: (i) subcutaneous adipose tissue, (ii) visceral adipose tissue, (iii) breast, (iv) skeletal muscle, (v) heart, (vi) thyroid, (vii) kidney, (viii) prostate, (ix) pancreas, (x) testis, (xi) liver, (xii) colon, (xiii) skin, (xiv) stomach and (Xv) lung, with majority cell types indicated in connected boxes. (**C**) Total number of expressed genes (in at least one tissue type) by respective status: (i) no cell type enrichment in any tissue, (ii) prediction as cell type enriched in one tissue, or (iii) predicted to be cell type enriched in two or more tissues. Gene ontology overrepresented terms for genes with: (**D**) no predicted cell type enrichment and (**E**) predicted enrichment only in testis. (**F**) Cell type enrichment predictions for genes classified as enriched in seven or more tissue types. See also Table S1 and S2 and Figure S3.

Of the 19,634 protein coding genes expressed in one or more tissues, 5644 (28.7%) were not predicted to be cell type enriched in any tissue (Figure 2 C.i). GO analysis identified the most significant over-represented pathways among these genes as ’*metabolism of RNA*’ (FDR 4.6 ×10^−21^), ’*gene expression (transcription)*’ (FDR 2.3 ×10^−11^) ’*RNA polymerase II transcription*’ (FDR 5.4 ×10^−10^) and ’*rRNA processing* ’ (FDR 5.8 ×10^−10^) (subgroups shown in Figure 2 D), consistent with housekeeping function. Indeed, 2893 of these 5644 genes (52.3%, p<10^−15^) had been previously categorised as members of the housekeeping proteome (Uhlen et al., 2015).

5979 (30.4%) genes were classified as cell type enriched in only a single tissue (Figure 2 C.ii), the largest proportion of which were in testis (n=3141) (Table S1, Tab 4). GO term analysis of this gene group identified the most significant over-represented pathways as ’*sexual reproduction*’ (FDR 3.7 ×10^−32^) and ’*spermatogenesis*’ (FDR 2.9 ×10^−30^) (subgroups shown in Figure 2 E). Of the 8011 genes predicted to be cell type enriched in multiple tissues (Figure 2 C.iii), a small number (741, 9.2%) were enriched in seven or more; the majority of which were predicted to be immune cell-, endothelial cell- or stromal cell-enriched (Figure 2 F), i.e., in cell types profiled in all, or most, tissues. Enrichment scores for all genes in cell types by tissue type can be found in Table S2 (summary of cell type gene enrichment across tissue in Table S1, Tab 4).

### *Ref.T*. analysis can predict source of tissue enriched genes

RNAseq data from unfractionated human tissues can be used to identify genes with higher expression in any given tissue, compared to others. For genes classified as *tissue enriched* in the Human Protein Atlas (HPA) (Uhlen et al., 2015), those we classified as cell type enriched were predominantly expressed in tissue specialised cell types, for example, heart enriched genes were predominantly cardiomyocyte enriched and liver-enriched genes predominantly hepatocyte enriched (Figure S3 A). A hypergeometric test was performed to determine similarity between predicted cell type enriched genes and the top 300 enriched genes in each tissue in the GTEx data (Consortium, 2015) (as collated in the Harminozome database (Rouillard et al., 2016)); similar to the comparison with the HPA data, the highest statistical overlap between tissue enriched genes and cell enriched genes were predominantly with tissue specialised cell types (Figure S3 Bi-vi). This highlights the usefulness of our analysis of bulk RNAseq to disentangle cell type variance across the different tissues in the human body, independent of scRNAseq data.

### Pancreatic alpha and beta cells have both specific and shared gene enrichment profiles

Alpha and beta cells, the most abundant endocrine cell types in the pancreatic islet of Langerhans (Moede et al., 2020), are defined by their expression of the blood glucose elevating or lowering hormones, glucagon (*GCG*) and insulin (*INS*), respectively. As a general rule, transcripts predicted to be cell type enriched generally separated well from others, but analysis of pancreas samples (n=328) revealed that many transcripts that correlated most highly with the alpha cell *Ref.T*. panel also correlated well with the beta cell *Ref.T*. panel (Figure 3 A.i), and *vice versa* (Figure 3 A.ii). Analysis of individual transcripts revealed 131 genes highly and selectively correlated with the *Ref.T*. panels for *both* alpha and beta-cells (Figure 3B, [grey central panel; mean differential corr. between Ref.T panels <0.15]). GO and reactome analysis (Ashburner et al., 2000) of these 131 co-enriched genes revealed over-represented classes and pathways included ’*regulation of secretion by cell*’ (FDR 7.5 ×10^−11^), ’*neuronal system*’ (FDR 9.9 ×10^−7^) and ’*synapse*’ (FDR 1.5 ×10^−15^) (Table S3, Tab 1, Tables A-C). Synapse related proteins (n=44) included members of the synaptotagmin (*SYT4, 5, 7, 13, 14*), and glutamate receptor (*GRIA2,3*) families (Table S3, Tab 2), many of which are reported to be important for pancreatic endocrine cell function e.g., *SYT4* (Huang et al., 2018) and *SYT13* (Bakhti et al., 2021; Tarquis-Medina et al., 2021), whilst the function of others in this context is not currently known e.g., *FRRS1L and NSG1*. Alpha and beta cell co-enriched genes included several encoding for transcription factors involved in islet cell specification, e.g., *NKX2-2*, (Churchill et al., 2017), *NEUROD1* (Bohuslavova et al., 2021), *RFX6* (Soyer et al., 2010), *INSM1* (Liang et al., 2021), *PAX6* (Hart et al., 2013) and *MYT1* (Wang et al., 2007), as well as those with no currently reported function in these cell types, e.g., *CELF3* and *MYT1L* one could speculate such genes likely have a role in neuroendocrine cell function. 91 genes had predicted alpha cell-enrichment, including *GCG, TTR* and *KCNH6* (Figure 3 B, left side); all of which are involved in glucose homeostasis (Noguchi and Huising, 2019; Su et al., 2012; Yang et al., 2018), and other genes with, as yet, no described function in this cell type e.g., *SMIM24, CALY* and *C5orf38* (Figure 3 B, left side). 69 genes had predicted beta cell enrichment, including those encoding proteins with known beta cell-specific functions, e.g., *IAPP* and *MAFA* (Nishimura et al., 2015; Westermark et al., 2011), as well as those with no reported function in this cell type, e.g., *HHATL, SNCB* and *SLC6A17* (Figure 3 B, right side). Tissue profiling for selected genes showed protein expression consistent with our classifications (Figure C-D top panel). We sourced data from scRNAseq of human pancreas (Tabula Sapiens et al., 2022), to compare the expression profiles of selected predicted alpha- (Figure 3 C.i-iv), beta- (Figure 3 E.i-iv) or co- (Figure 3 D.i-iv) enriched genes; categorisation was largely consistent between datasets. A small number of genes we predicted to be alpha-, beta or co-enriched had a mean expression <0.1 TPM in the analysed bulk RNAseq dataset (gene n=11, 6 and 4, respectively, Figure S4 A). Despite this low expression, our predicted expression of these genes was consistent with the scRNAseq analysis; with most (21/22 [95%]) detected predominantly in the correspondingly annotated cell types (Figure S4 C-E). However, for several of these genes, detectable expression by scRNAseq was low, or only evident in a small number of cells within the cluster, e.g., *GLB1L3* (Figure S4 C.ii). The interpretation of such scRNAseq data is challenging; thus, our classifications, based on a completely independent data collection and analysis method, can be used to verify that low or irregular detection of gene expression by scRNAseq in an annotated cell type supports a real biological phenomenon, as opposed to noise or imputation artefact.

**Figure 3.**
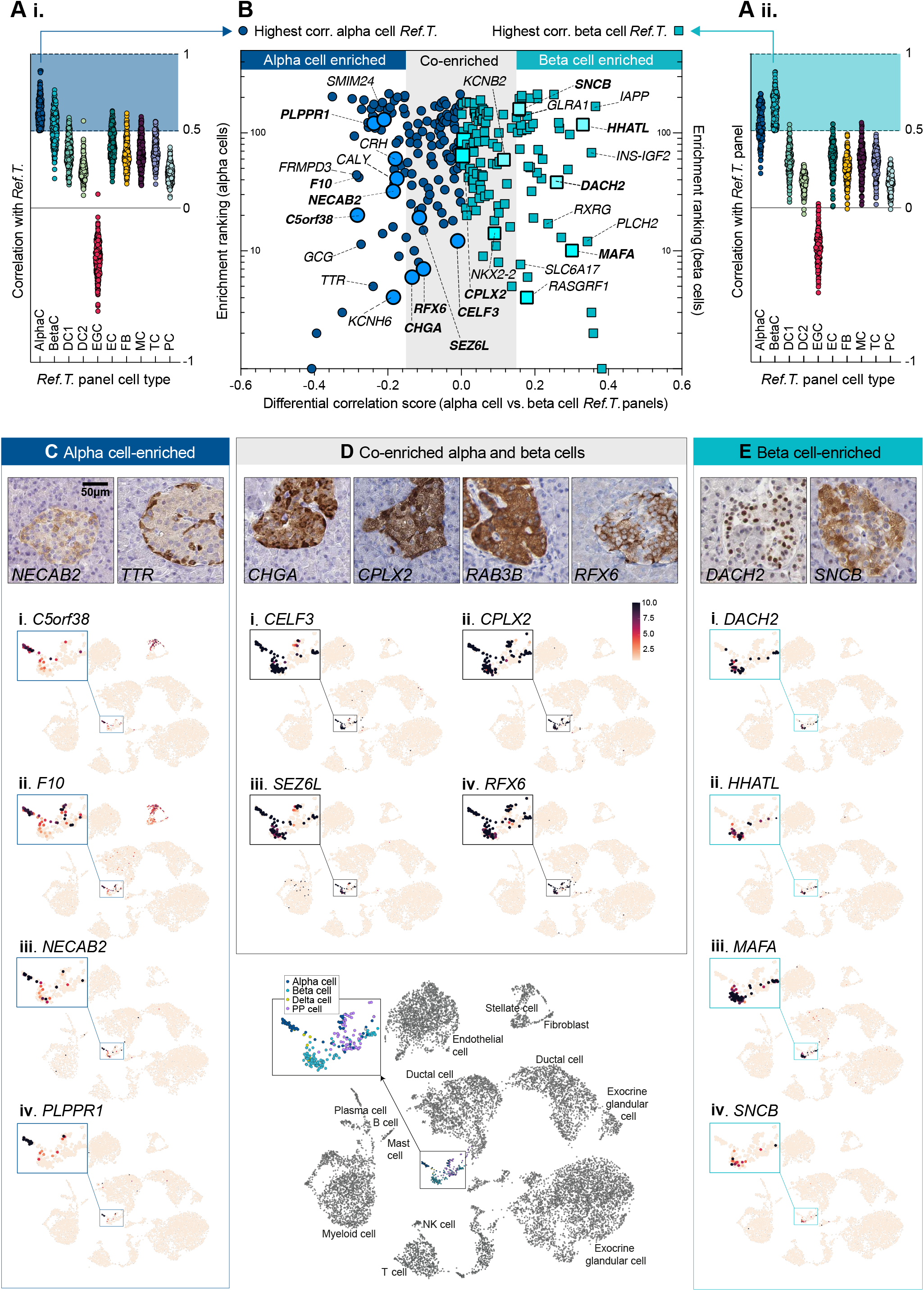
Pancreatic alpha and beta cells express respective cell type enriched genes and a panel of shared co-enriched genes. RNAseq datasets for human pancreas (n=328) were retrieved from GTEx V8 and correlation coefficients between selected cell type *Ref T*. and all others were generated. Mean correlation values between protein coding genes that correlated most highly with (i) alpha or (ii) beta cell .*Ref.T*. (above >0.50) and all *Ref.T*. panels. (**B**) For these transcripts, the ‘*differential correlation score’(differ*ence between mean correlation with alpha and beta cell *Ref.T*.) was plotted vs. ‘enrichment ranking’(position in each respective list, highest correlation = rank 1). Shaded grey box highlights genes with differential correlation <0.15. Genes highlighted in bold correspond to those featured in the lower panels. Tissue protein profiling of selected genes predicted to be (**C**) alpha cell-enriched, (**D**) co-enriched in both alpha and beta cells, or (**E**) beta cell-enriched, in human pancreas samples. scRNAseq data from analysis of human pancreas was sourced from Tabula Sapiens (Tabula Sapiens et al., 2022), and used to generate UMAP plots, showing the expression profiles of example genes we predicted as being (**C**) alpha cell-enriched; (i) *C5orf58*, (ii) *F10* (iii) *NECAB2* and (iv) P*LPPR1*, (**D**) co-enriched in both alpha and beta cells; (i) CE*LF3*, (ii) *CPLX2*, (iii) *SEZ6L* and (iv) *RFX6*, or (**E**) beta cell-enriched; (i) *DACH2*, (ii) *HHATL*, (iii) *MAFA* and (iv) *SNCB*. scRNAseq cell type annotations are displayed on lower central plot. AlphaC; alpha cell, betaC; beta cell, DC1; ductal cell 1, DC2; ductal cell 2, EC; endothelial; FB1/2; fibroblast 1/2, SMC; smooth muscle cell, MC; macrophage, MastC; mast cell, NP1/2; neutrophil 1/2, TC; T-cell, PC; plasma cell. see also Table S3 and Figure S4.

### Temporal changes in gene enrichment signature underlie the process of spermatogenesis

Precise definitions, markers and terminology used for the respective cell types in the different stages of spermatogenesis vary between studies. Our analysis was based on four *Ref.T*. panels (S1-S4) that were selected to represent the temporal order of development: S1, germ cell expressed [*MAGEB2, KDM1B, PIWIL4*] (spermatogonia), S2, meiotic cell cycle expressed [*ANKRD31, RBM44, TOP2A*] (spermatocytes), S3, spermatid structure-related [*CEP55, KPNA5, PBK*] (round/early elongating spermatids) and S4, nuclear condensation/protamine repackaging factors [*PRM1, PRM2, TNP1*] (late/elongated spermatids) (Figure 4 A and Table S1, Tab 1, Table N). When the sample set was analysed by WGNCA, *Ref.T*. within each respective panel were all in a common module (Figure S5 A). Principle component analysis of the corr. values of cell-enriched genes vs. all *Ref.T*. panels revealed the greatest proportion of variance in enrichment, and thus uniqueness vs. other cell types, was driven by cell types S1, S2, S3, S4 (Figure 4 B). Tissue profiling for proteins encoded by a panel of genes predicted to be enriched in cell types outside those in the spermatogenesis pathway revealed expression consistent with our classifications (Figure 4 C). 6179 genes were enriched in one or more of the germ cell types representing the different stages of spermatogenesis, vs. non-germ cell types (Figure S5 B.i and Table S4, Tab 1 [correlation with respective Ref.T panel >0.50, differential correlation vs. all non-germ cell types >0.15] columns H-K and Q). GO and reactome analysis of this gene list revealed that the most significantly over-represented terms included ’*sexual reproduction*’ (FDR 3.1 ×10^−27^), ’*microtubule-based processes*’ (FDR 2.2 ×10^−26^), ’*male gamete generation*’ (FDR 2.3 ×10^−26^) and ’*cell cycle*’ (FDR 4.6 ×10^−19^) (Table S4, Tab 2, Tables A and B) (Figure S5 B.ii [summary plot of GO terms, made with REVIGO (Supek et al., 2011)]). Genes that correlated with *Ref.T*. panels representing cells at different stages of spermatogenesis had two main profiles; they were enriched at a specific developmental stage, i.e., S1 (Figure 4 D.i), S2 (Figure 4 D.ii) S3 (Figure 4 D.iii) or S4 (Figure 4 D.iv) (for all see Figure S5 C .i and ii) or, they were co-enriched in adjacent cell types on the developmental trajectory: i.e., S1 and S2 (Figure 4 D.v), S2 and S3 (Figure 4 D.vi), S3 and S4 (Figure 4 D.vii) or S2, S3 and S4 (Figure 4 D.viii) (for all see Figure S5 D.i and ii). Each plot shows five illustrative genes for each enrichment profile type (Figure 5 Ei-vii), including genes encoding for proteins with a previously reported function at the corresponding stage of spermatogenesis e.g., for S1: *FGFR3* (Ewen et al., 2013), and those with no known role in this context e.g., *C19orf84*(Figure 4 E.i). Protein profiling revealed spatial distribution for those encoded by genes classified as S1, S2, S3 or S4-enriched or co-enriched, with positive signals observed progressively closer to the centre of the seminiferous tubule with each subsequent developmental stage (Figure 4 E.i-vi). GO analysis revealed over-represented classes in genes predicted to be S1, S2- or S1 & S2 enriched included developmental, cell cycle and meiotic processes (Figure 4 F.i, ii and v), whilst organelle assembly, microtubule processes and cilium and flagellum organisation and motility associated genes were over represented in S3-, S4- and S3 & S4-enriched genes (Figure 4 F.iii, iv and vii) (Table S4, Tab 3). No transcripts were predicted to be co-enriched in non-adjacent cell stages along the developmental trajectory (e.g., S1 and S3, or S2 and S4), consistent with a coordinated temporal modification in gene enrichment signatures between subsequent stages. A single gene, *MEIOC*, was predicted to be enriched in 3 stages - S2, S3 and S4. *MEIOC* is required for germ cells to properly transition to a meiotic cell cycle program, together with binding partners *YTHDC2* and *RBM46* (Qian et al., 2022); both of which we also predicted as enriched in cells in S2 and, to a lesser extent S3 (Table S4, Tab 1). Data from scRNAseq of human adult testis (Guo et al., 2018) supported our predictions, showing *MEIOC* enrichment in cell clusters broadly corresponding to our classification of S2, S3 and S4 (Figure 4 G.i) (cell type annotation UMAP as in the original publication in Figure S5). In contrast, we predicted that the related transcript *MEIOB* had specific enrichment at stage S2 (Figure 4 D.ii and E.ii), which was also verified by scRNAseq (Figure 4 G.ii). scRNAseq for selected less well characterised genes that we predicted as enriched in either S1, S2, S3 or S4 cells (Figure S5 C.iii), or gene predicted to be co-enriched in two stages (Figure S5 D.iii) also showed agreement with our classifications. A number of genes that were predicted to be enriched in one or more of the germ cell stages were lowly expressed (n=240 with mean TPM<0.1), several of which did not appear in the scRNAseq dataset (Guo et al., 2018), presumably due to a lack of detection. Of the 100 most lowly expressed genes for which scRNAseq data was available, most (>80%) had expression profiles consistent with our predictions in the scRNAseq data (examples in Figure S6), but in many cases transcripts were detected at low levels in only a few cells in the corresponding cluster, e.g., *FZD10* (Figure S6 D), *LEP* (Figure S6 F) and *SIGLEC15* (Figure S6 J), making interpretation of this scRNAseq data in isolation challenging. Thus, we show that analysis of bulk RNAseq can identify differentially enriched genes associated with one or multiple stages of the developmental trajectory during spermatogenesis, including genes that are likely too lowly expressed for detection or classification as cell type enriched by scRNAseq.

**Figure 4.**
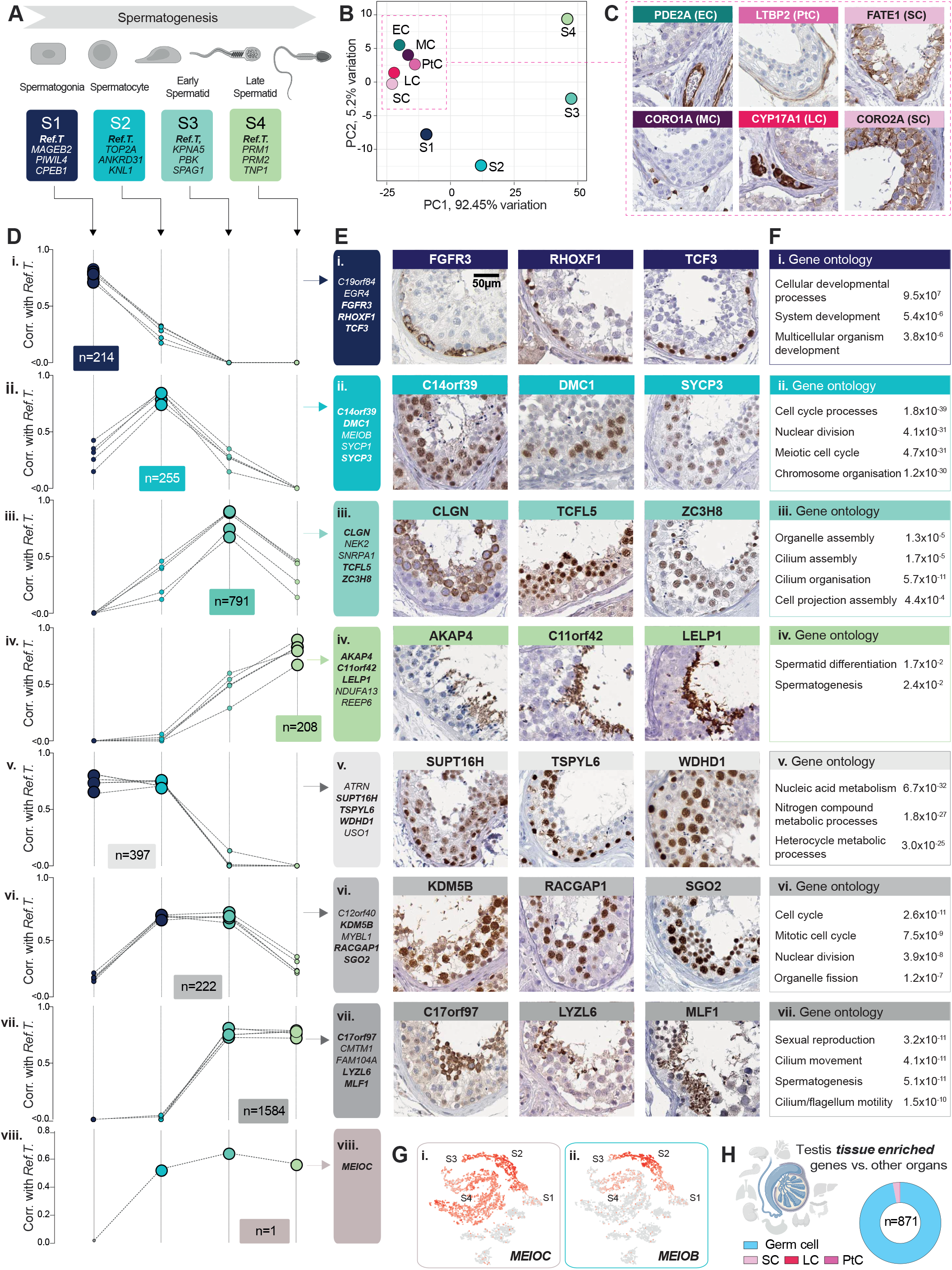
Analysis of pseudo temporal changes during spermatogenesis reveals stage-specific and common stage-shared gene enrichment signatures. (**A**) Cell types at the different stages of spermatogenesis were defined based on *Ref.T*. selected to broadly represent the developmental stage classifications spermatogonia [S1], spermatocytes [S2], early spermatids [S3] and late spermatids [S4]. **(B)** Principal component analysis of comparative correlation profiles of cell-enriched genes in S1, S2, S3, S4, sertoli cells (SC), Leydig cells (LC), peritubular cells (PtC), endothelial cells (EC) or macrophages (MC) vs. all *Ref.T*. panels. (**C**) Tissue profiling for proteins encoded by example genes we predicted to be enriched in non-germ cell types. (**D**) Pseudo trajectories of gene enrichment signatures over time, showing enrichment values for each developmental stage, using (**E**) illustrative genes predicted to be: (i) S1, (ii) i) S3, (iv) S4, (v) S1 and S2, (iii)S3, (vi) S2 and S3, (vii) S2, S3 and S4, enriched, with corresponding tissue protein profiling. (**F**) Over-represented gene ontology terms and significance corrected FDR values for all genes classified as: (i-iv) highly enriched at a specific stage, or (v-vii) co-enriched at one or more stages of development. (**G**) UMAP plots showing gene expression profile in the Human Testis Atlas scRNAseq data (Guo et al., 2018) of: (i) the S2, S3 and S4 predicted enriched gene *MEIOC* and (ii) the S2 predicted enriched gene *MEIOB*. (**H**) Classification of testis tissue enriched genes that we predicted to be cell type enriched. See also Table S4, Figure S5 and S6.

**Figure 5.**
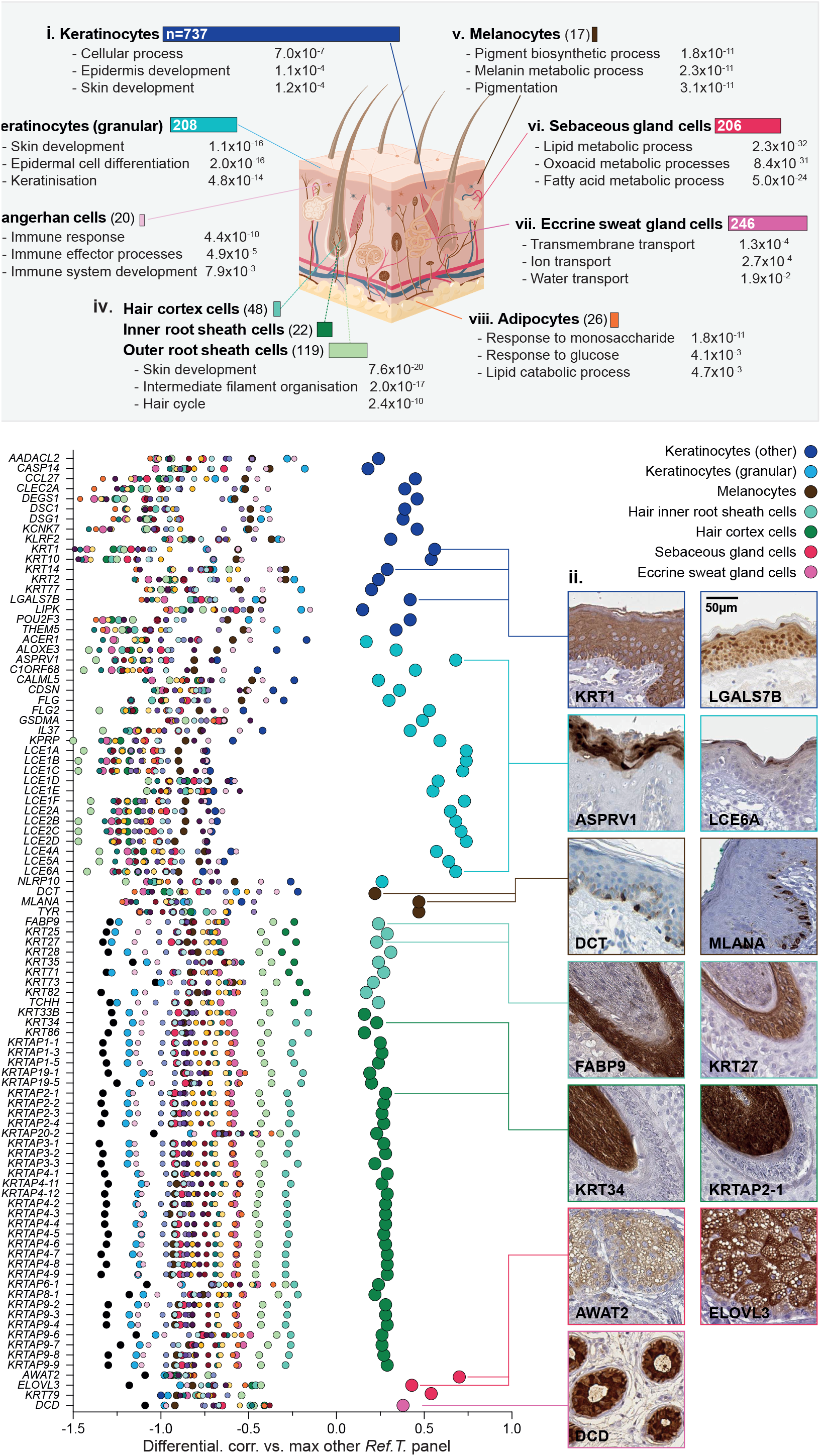
Constituent cells of the skin hair root are the primary source of skin tissue enriched genes. (**A**) Number of genes predicted to be cell type enriched, and corresponding over-represented gene ontology terms and significance corrected FDR values, for skin specialised cell types profiled: (i) keratinocytes, (ii) granular keratinocytes, (iii) Langerhans cells, (iv) hair cortex cells, inner and outer root sheath cells, (v) melanocytes, (vi) sebaceous gland cells (vii) eccrine sweat gland cells and (viii) adipocytes. (**B**) (i) Genes enriched in skin vs. other organs, which were predicted to be cell type enriched in our analysis, were plotted to show the min differential values between the mean correlation coefficients with the *Ref.T*. panels for each cell type. Enlarged circles represent classification as predicted cell type enriched. (**ii**) Tissue profiling for proteins encoded by skin tissue enriched genes with predicted enrichment in the indicted cell type. See also Figure S7.

RNAseq data from unfractionated tissue can be used to identify genes with enriched expression in testis vs. other tissues, as we previously described (Uhlen et al., 2015). The vast majority of genes with testis-enriched expression were predicted to be enriched in one or more germ cell type (845/871 [97%]), with a smaller number predicted to be enriched in sertoli (24/871 [2.8%]), Leydig (24/871 [0.1%]) or peritubular cells (24/871 [0.1%]) (Figure 4H). No testis enriched genes were classified as endothelial or macrophage-enriched in our analysis, reflecting their presence in other tissues.

### Constituent cells of the skin hair root are the primary source of skin tissue enriched genes

The skin is one of the most complex tissue types in the human, with multiple layers and diverse constituent cell types. We profiled 18 different cell types in the skin, many of which are not represented in scRNAseq data in Tabula Sapiens (Tabula Sapiens et al., 2022) or the Human Protein Atlas (HPA) (www.proteinatlas.org/) (Karlsson et al., 2021; Uhlen et al., 2015), e.g. sebaceous gland cells, eccrine sweat gland cells, adipocytes, hair cortex and inner/outer root cells. Keratinocytes expressed the largest proportion of predicted cell type-enriched genes; 737 in the sub-granular layers (Figure 5 A.i) and 208 in the granular layer (Figure 5 A.ii). Analysis of the sub-granular keratinocyte layers at a higher cell type resolution was not possible, as a *Ref.T* panel with high, consistent, specificity for sub-population(s) of basal and suprabasal keratinocytes could not be identified. Similarly, when the dataset was analysed by WGNCA, most genes we predicted to be sub-granular keratinocyte enriched clustered in multiple groups on common clades (552/737 [75%]), the constituent groups of which contained a combination of genes considered basal e.g., *COL17A1* or suprabasal e.g., *DSG1* keratinocyte markers (Figure S7 A.i). In contrast, *Ref.T*. representing granular keratinocytes and the majority of genes predicted to be enriched in this cell type (181/208 [87%]), clustered in two groups on a single clade (Figure S7 A.ii). These results are consistent with keratinocyte development being associated with a shift in absolute gene expression levels, as opposed to a defined transition between highly distinct cell states that express many specific markers (prior to terminal differentiation in the granular layer).

For genes identified as cell type enriched, GO analysis revealed over-represented classes consistent with cell type annotation, e.g. for granular keratinocytes significant terms included ’*epidermal cell differentiation*’ (FDR 2.0 ×10^−16^) (Figure 5 A.ii) and for sebaceous gland cells ’*lipid metabolic processes*’ (FDR 2.3 ×10^−32^) (Figure 5 A.vi). Of the skin-specific cell types profiled, melanocytes had the fewest enriched genes (n=17) (Figure 5 A.v), including highly expressed genes with known cell type-specific functions e.g., *PMEL, DCT* (mean TPM in skin RNAseq 58.2 and 29.6, respectively). More lowly expressed melanocyte-enriched genes included *SLC24A5, CA14* and *SLC45A* (mean TPM in skin RNAseq 0.5, 1.9 and 5.7, respectively). In skin scRNAseq data from Tabula Sapiens (Tabula Sapiens et al., 2022) (Figure S7 B.i) *SLC24A5* was predominantly expressed in a sub-population of cells in melanocyte annotated cluster (Figure S7 B.ii), but *CA14* and *SLC45A2* were not as clearly enriched in this cell type (Figure S7 B.iii and iv). However, our classifications of these genes as melanocyte-enriched are supported by other studies showing that *SLC45A2* has a role in deacidification of maturing melanosomes to support melanin synthesis (Le et al., 2020) and that *CA14* is downregulated in vitiligo skin samples, compared to normal, along with other genes we classified as melanocyte enriched (Yu et al., 2012). Furthermore, all three of these genes were clustered together with the melanocyte *Ref.T* when the dataset was analysed by WGNCA (Figure S7 A.iii). Thus, as we demonstrated for alpha and beta cells in the pancreas and germ cells in the testis, our analysis can identify cell-type enriched genes that are not always detectable as such by scRNAseq.

RNAseq data from unfractionated tissue was used to identify 188 genes as skin enriched vs. other tissues in the HPA tissue section (Uhlen et al., 2015), of which 151 were also identified as such in a similar analysis of tissues in GTEx (Consortium, 2015), collated in the Harminozome database (Rouillard et al., 2016). Of these, 96/151 [63%] were predicted to be cell type enriched in our analysis (Figure 5 B.i); most frequently in cells of the hair root (hair cortex or inner root sheath cell), granular keratinocytes or other keratinocytes. Other skin enriched genes were predicted to be enriched in melanocytes, sweat gland or sebaceous gland cells (Figure 5 B.i). Tissue profiling of proteins encoded by selected genes supported our classifications (Figure 5 B.ii). No skin enriched genes were predicted to be cell type enriched in endothelial cells, smooth muscle cells, fibroblasts, macrophages, or other immune cell types - consistent with their presence in multiple tissue beds, and thus lack of specificality to skin. Of those cells that were skin enriched, but not classified as cell type enriched in our analysis (Figure S7 C.i [rows lacking an enlarged circle]) most had co-enrichment profiles in multiple cell types in the hair root (Figure S7 C.ii). These genes included *PSORS1C2*, a member of the psoriasis susceptibility locus (Abbas Zadeh et al., 2017). Enrichment of this gene in cell types of the hair follicle is supported by studies showing that ’near naked hairless’ mice, which have a spontaneous mutation preventing the development of a normal coat, have significantly reduced expression of *PSORS1C2* (Liu et al., 2007), together with others highlighted here e.g., *S100A3* and *KRTAP16-1* (Figure S7 C.ii) (Liu et al., 2007). In depth skin tissue profiling showed expression of selected encoded proteins consistent with enrichment in the hair root (Figure S7 C.iii). Previously, keratinocytes, the majority cell type in the skin, have been annotated as the site of expression for the majority of skin enriched genes (Karlsson et al., 2021). However, this is likely due to the lack of hair root cells in the scRNAseq data on which these classifications are based. Here, we show that a minority cell type represents the most common source of skin enriched genes.

### Cross-tissue analysis reveals similarities in cell type gene enrichment signatures

8011 genes were predicted to be cell type enriched in more than two tissue types. To explore the relationship between these cell type gene enrichment signatures, we performed a hypergeometric test to determine the degree of similarity between all cell types in all tissues. As cell type gene enrichment signatures are generated via a correlation-based analysis, independent of cell type absolute gene expression levels, such comparisons between tissue datasets can be made without correction for batch effects, the analysis platform used, or requirement for normalisation.

### Organ-specific cell types can have common gene enrichment signature panels

Organ specific cell types (i.e. excluding those found in all or multiple organs, e.g., endothelial cells, fibroblasts and immune cell types) had gene enrichment signatures with: little or no similarity to other cell types e.g., hair inner root sheath cells and melanocytes (Figure 6 A.ii and iii), significant similarity to one other cell type, e.g., skeletal myocytes and cardiomyocytes (Figure 6 A.iv) or significant similarity with multiple other cell types, e.g., endocrine cells from several tissues; alpha and beta cells from the pancreas, enteroendocrine cells from the colon and stomach, and parafollicular cells from the thyroid (Figure 6 A.vi). We found a significant overlap between the enriched gene signatures of adipocytes (subcutaneous adipose, visceral adipose and breast), skin sebaceous gland cells, liver hepatocytes, and kidney proximal tubular cells (Figure 6 A.vii). GO analysis of the 41 genes predicted to be enriched in at least three of the aforementioned cell types (Figure 6 B.i [green box]) (Table S5, Tab 1, Table A) revealed significant terms all related to metabolic processes, including ’*carboxylic acid metabolic processes*’ (FDR 8.8 ×10^−26^) and ’*organic acid metabolic processes*’ (FDR 9.4 ×10^−26^) (Table S5, Tab 1, Table B) (Figure 6 B.ii). 22 of these 41 genes were also predicted to be enriched in cardiomyocytes, another highly metabolically active cell type with a significant overlap in gene enrichment signature with both adipocytes and hepatocytes (Figure 6 A and Table S5, Tab 1, Table A). Illustrative protein profiling showed selective expression of ACO1 and HADH in adipocytes in adipose tissue, sebaceous gland cells in skin, hepatocytes in liver and proximal tubular cells in kidney (Figure 6 B.iii). The enrichment of such genes in many highly metabolically active cells is consistent with a common shared function across tissue types. In contrast, cell type enriched genes classified as such in only one tissue are likely key for highly specialised cell functions, e.g., complement and coagulation factor genes were predicted to be enriched only in hepatocytes (including *C4B, C8A, C9, CFHR1/2/4/5* and others) and specific solute transporters only in proximal tubular cells (e.g., *SLC13A1, SLC22A13, SLC22A6, SLC22A8*). Tissue profiling for proteins encoded by example genes predicted to be enriched in only one of these four cell types; adipocytes (*PRKAR2B*), sebaceous gland cells (*TMEM97*), hepatocytes (*OTC*) or proximal tubular cells (*TMEM174*) showed positive staining in only the respective cell types (Figure 6 B.iv). In contrast to ACO1 and HADH, which were expressed mean TMP >10 in the RNAseq datasets analysed (Figure 6 B.v), expression values of these example genes were highest in the corresponding tissue, with low or no expression in the others (Figure 6 B.vi).

**Figure 6.**
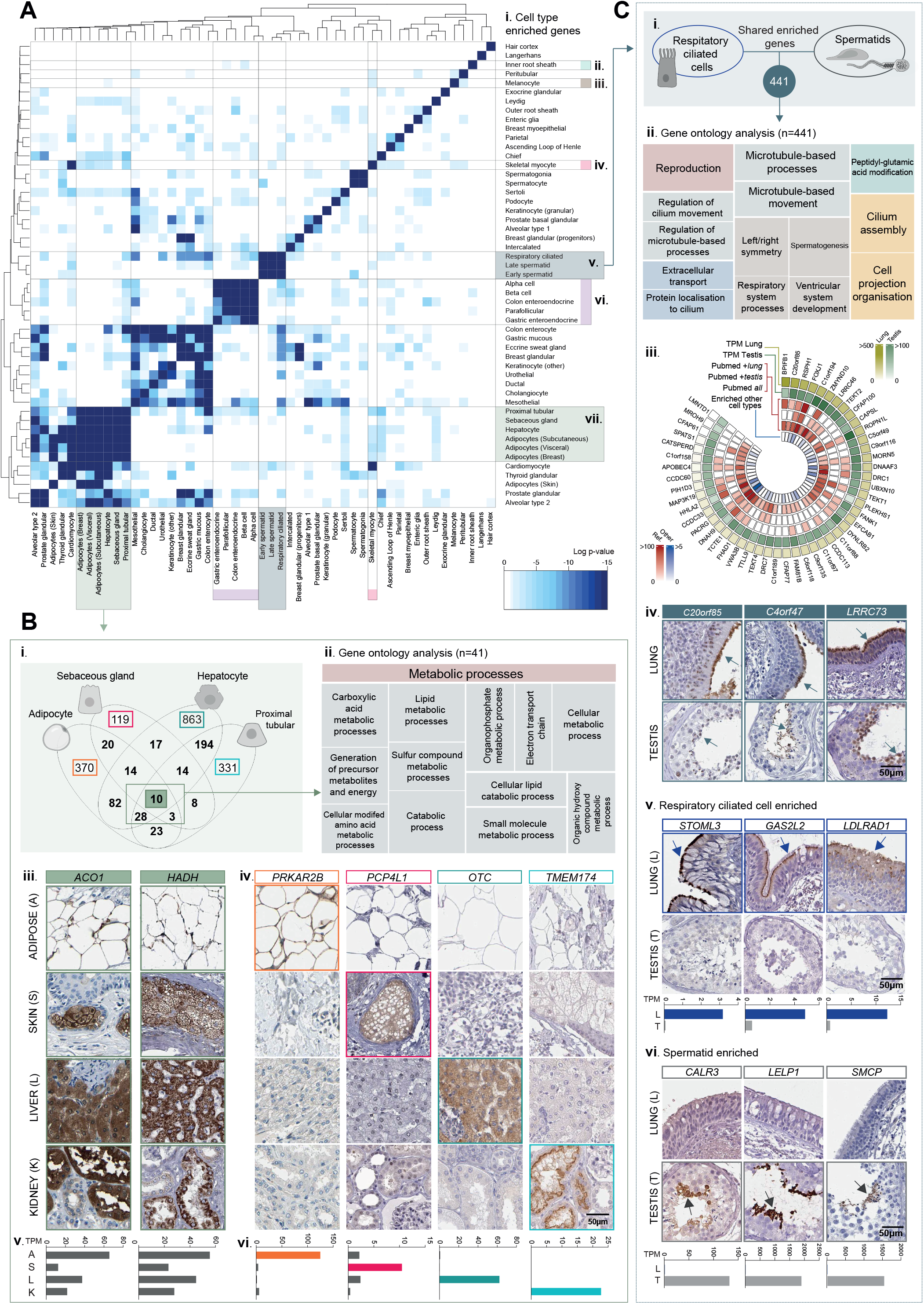
Organ specific cell types can have gene enrichment signature similarities. (**A**) Heatmap showing significance p-values for similarity scores between predicted cell type enriched genes, calculated using a hypergeometric test for: (i) all organ-specific cell type enriched gene signatures, (ii) skin inner root sheath hair cells, (iii) skin melanocytes, (iv) skeletal myocytes, (v) lung respiratory ciliated cells and testis spermatids, (iv) pancreatic alpha and beta cells, colon enteroendocrine cells, thyroid parafollicular and stomach gastric enteroendocrine cells and (vii) kidney proximal tubular cells, sebaceous gland cells, hepatocytes, and adipocytes in subcutaneous adipose tissue, visceral adipose tissue and breast. (**B**) (i) Number of individual or common enriched genes for adipocytes (in at least 2/3 tissues profiled) kidney proximal tubular cells, sebaceous gland cells and hepatocytes. (ii) Over-represented gene ontology terms among the 41 genes featuring in the gene enrichment signature of at least 3/4 cell types, displayed in tree map format, generated using REVIGO (areas proportional to Log10 significance values). Tissue profiling for proteins encoded by genes that were: (iii) part of the shared gene enrichment signature [ACO1, HADH] or (iv) classified as cell type enriched only in one cell type [PRKAR2B, TMEM97, OTC, TMEM174] and corresponding mean TMP expression in the corresponding tissue RNAseq datasets (v) and (vi), respectively. (**C**) (i) Number of genes that featured in gene enrichment signatures for respiratory ciliated cells, early and late spermatids and (ii) the over-represented gene ontology terms among these shared enriched genes. (iii) Circular plot showing up to the top 50 most enriched genes in respiratory ciliated cells and spermatids, displaying the mean TMP values in the lung and testis RNAseq datasets, the number of mentions in Pubmed of gene *and* corresponding tissue (’*Pubmed + lung/testis*’) or the gene alone (’Pub*med all*’), and the number of other cell types in which the gene was also predicted to be enriched (’enri*ched other cell type*’). Tissue profiling for proteins encoded by genes with predicted enrichment in:(iv) both respiratory ciliated cells and spermatids, (v) respiratory ciliated cells only and (vi) spermatids only and corresponding mean TMP values in the tissue RNAseq. See also Table S5 and Figure S8.

Our analysis also revealed a significant overlap between the gene enrichment signatures of early and late spermatids in the testis and respiratory ciliated cells in the lung (Figure 6 A.v and C.i). GO term analysis of these 441 shared genes (Table S5, Tab 2, column A-B) revealed the most significant terms were related to cilia (Figure 6 C.ii), which are important for both clearance of fluid from the airways and movement of the sperm flagellum, including ’*cilium organisation*’ (FDR 3.6 ×10^−69^) and ’*cilium movement*’ (FDR 9.5 ×10^−64^) (Table S5, Tab 2, Table 1). The top 50 genes predicted to be most highly enriched in both early and late spermatids and RCC (Figure 6 C.iii) had variable absolute expression in the respective tissues. *LMNTD1* and *MROH9* had very low expression in the lung RNAseq (mean TMP 0.42 and 0.68, respectively) (Figure 6 C.iii) and scRNAseq data from human lung (Tabula Sapiens et al., 2022) revealed highly specific, but variable expression (or detection) of these genes in RCC (Figure S8 A.ii and iii). Predicted expression in S3 and S4 cells in testis was also supported by scRNAseq from the Human Testis Atlas (Guo et al., 2018) (Figure S8 B.ii and iii). Despite the highly specific enrichment profiles of *LMNTD1* and *MROH9*; neither were predicted to be enriched in any other cell type across all tissues analysed (Figure 6 C.iii), there are no existing studies of these genes in this context. Some other genes with highly predicted enrichment in early and late spermatids and RCC were also predicted to be enriched in several other cell types e.g., *PACRG* (Figure 6 C.iii), which is well studied in the context of motile cilia, particularly in sperm (Li et al., 2015), but has also been proposed to have other roles, such as in primary cilia (Dawe et al., 2005) and even inflammatory pathway signalling (Meschede et al., 2020); perhaps explaining its more widespread enrichment profiles in our analysis. Tissue profiling for proteins encoded by example genes enriched in both RCC and early and late spermatids (Figure 6 C.iv), or RCC only (Figure 6 C.v) showed expression consistent with our predictions. GO analysis of genes predicted to be highly enriched in spermatids, but not RCC, revealed the most significant terms were unrelated to cilia formation, including ’*spermatogenesis*’ (FDR 1. ×10^−25^), ’*multicellular organism reproduction*’ (FDR 1.5 ×10^− 19^), ’*spermatid development*’ (FDR 6.7 ×10^−14^) and ’*fertilisation’* (FDR 1.4 ×10^−10^) (Table S5, Tab 3, Column A-B and Table 1); reflecting an enrichment for genes with highly specialised function within the testis only, e.g., *CALR3, LELP1* and *SMCP* (Figure 6 C.vi).

### Core cell types have common gene enrichment signature panels across tissues

Eight cell types were profiled in all, or most, tissue types (termed “core cell types”); endothelial cells [n=15 tissues], smooth muscle cells [n=10], fibroblasts (including hepatic stellate cells (HSC) in the liver, and adipose progenitor cells (APC) in adipose tissue) [n=14], macrophages (including Kupffer cells in the liver) [n=15], neutrophils [n=8], mast cells [n=5], T-cells [n=13] and plasma cells [n=14] (Figure 7 A.i). Gene enrichment signatures of the same core cell type in different tissues had high similarity, with little, or no, crossover between different cell types (Figure 7A). Notable exceptions included hepatic stellate cells (HSC) and fibroblasts in liver and kidney, respectively, which had some commonality with smooth muscle cell gene enrichment signatures in other tissues (Figure 7 A.ii), in line with reports that liver HSC can have contractile properties (Soon and Yee, 2008) and potentially reflecting the presence of a kidney myofibroblast-like population, and lung neutrophils, which had some similarity to macrophages in several other tissues (Figure 7 A.iii). Enrichment signatures of core cell types had little or no cross over with those of organ specific cell types (Figure S8 C.i), except for lung macrophages, which had a significant similarity with the cell type group we previously identified as having shared gene enrichment signatures related to metabolic processes (Figure 6 B), including adipocytes, hepatocytes, proximal tubular cells (Figure S8 C.ii). One could speculate that this indicates macrophages in the lung have specific metabolic characteristics, in keeping with recent studies indicating that their metabolic responses to infectious pathogens or other insults may be distinct from other macrophage subtypes (Khaing and Summer, 2020).

**Figure 7.**
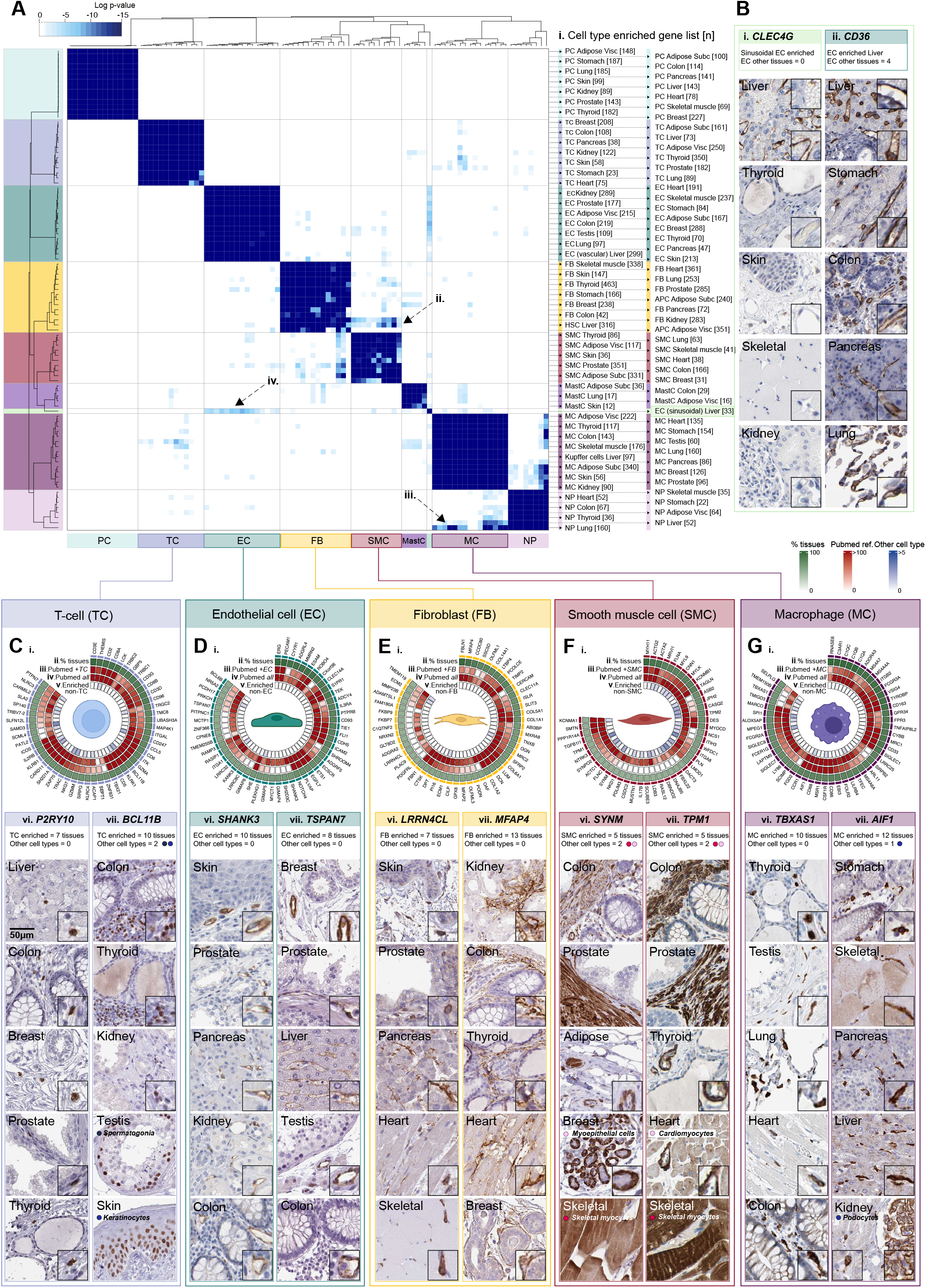
Core cell types share gene enrichment signatures across organs. (**A**) Heatmap showing significance p-values for similarity scores in cell type gene enrichment signatures, calculated using a hypergeometric test, between (**i**) plasma cells (PC), T-cells (TC), endothelial cells (EC), fibroblasts (FB), smooth muscle cells (SMC), mast cells (MastC), macrophages (MC) and neutrophils (NP) in different tissues. (**B**) Tissue profiling for proteins encoded by (i) the sinusoidal EC enriched gene *CLEC4G* and (ii) the vascular and sinusoidal EC enriched gene *CD36*, in different tissue beds. Circular plots showing up to the top 50 genes most frequently predicted as enriched in (**C**) TC, (**D**) EC, (**E**) FB, (**F**) SMC and (**G)** MC in different organs, displaying (i) the percentage of tissues in which the gene was classified as enriched in the given cell type (’*% tissues*’), the number of mentions in Pubmed of (ii) gene *and* corresponding core cell type (’*Pubmed + cell type*’) together or (iii) the gene alone (’*Pubmed all*’), and (iv) the number of other cell types (including non-core cell types) in which the gene was also predicted as enriched (’*enriched non-cell type*’). (v-vi) Tissue profiling of proteins encoded by selected genes predicted to be core cell type enriched. See also Table S6 and Figure S8.

Endothelial cells had strong gene enrichment signature similarities across tissues (Figure 7 A), with the exception of liver sinusoidal endothelial cells (LSEC), where over half of the enriched genes (19/34 [56%]) were not enriched in endothelial cells in any other tissue, consistent with their unique structural and phenotypic features, and highly specialised function (Shetty et al., 2018). Despite this, overall, they did have greatest similarity with vascular endothelial cells vs. any other core cell type (Figure 7 A.iv). Tissue profiling for proteins encoded by *CLEC4G* (Figure 7 B.i) and *CD36* (Figure 7 B.ii) showed expression consistent with our predictions of LSEC enrichment only, or vascular endothelial and LSEC enrichment, respectively.

To define key components of the gene enrichment signature for each core cell type, we identified genes predicted to enriched in at least half of the tissues profiled (Table S6), e.g., in at least 8/15 tissues for EC and MC (Figure 7 C-Gi and ii and Table S6, Tab 1). To assess existing reports for each gene in each given cell type, we used PubMed to search for the number of studies citing both gene name and cell type together (Figure 7C-G.iii), or gene name alone (Figure 7 C-G.iv). Many were well characterised genes on which a plethora of studies have been performed, e.g., *CD3E* and *CD2* in T-cells (Figure 7 C.iii and iv), others were poorly studied in the cell type context. For example, *SHANK3*, predicted to be endothelial cell enriched in 10 tissues (Figure 7 D), has been researched predominantly in the context of neurons and autism (Delling and Boeckers, 2021). *TSPAN7* (Figure 7D), predicted to be endothelial cell enriched in 8 tissues, has only been identified in endothelial cells in the context of tumour associated vasculature and metastasis (Sawada et al., 2022). There is little information in the literature about the function of *LRRN4CL* (Figure 7 E), a gene we predicted to be fibroblast enriched in 7 tissues, except for its elevated expression in skin melanoma metastases and breast cancer samples (van der Weyden et al., 2021; Zhang et al., 2021). In contrast, *MFAP4* (Figure 7 E) is a well-known gene in this cell context (Lin et al., 2020). *TBXAS1* was identified as a macrophage core enriched gene, and its enzymatic product, thromboxane A2, is linked to vasoconstriction and platelet aggregation, with links to innate immunity (Sellers and Stallone, 2008), but little knowledge exists in the macrophage context. Tissue profiling for proteins encoded by predicted endothelial enriched genes *SHANK3* (Figure 7 D.v) and *TSPAN7* (Figure 7 D.vi), fibroblast enriched genes *LRRN4CL* (Figure 7 E.v) and *MFAP4* (Figure 7 E.vi) and the macrophage enriched gene *TBXAS1* (Figure 7 G.vi), revealed selective expression consistent with our predictions.

Whereas most core endothelial and fibroblast enriched genes were not predicted to be enriched in any other cell types in our analysis (Figure 7 D-E.v), several T-cell (Figure 7 C.v) smooth muscle cell (Figure 7 F.v), and macrophage (Figure G.v) enriched genes where predicted to be enriched in an additional cell type(s). *BCL11B*, a gene we predicted to be T-cell enriched gene in 10 tissues (Figure 7 C) with a known role in T-cell development (Hosokawa et al., 2020), was also predicted to be enriched in skin keratinocytes, consistent with its role in dermal development in mice (Golonzhka et al., 2009), and unexpectedly, in the absence of any existing reports, in spermatogonia in the testis; expression profiles we verified by protein profiling (Figure 7 C.vii). *AIF1*, predicted to be macrophage enriched in 12 tissues (Figure 7 G.i and viii), consistent with its known expression in this cell type (Zhao et al., 2013), was also classified as kidney podocyte enriched; previously reported in only a single study (Tsubata et al., 2006), a prediction we again verified with tissue protein profiling (Figure 7 G.vi). Of all the core cell types, smooth muscle cell enriched genes were most likely to have predicted enrichment in another cell type, most frequently cardiomyocytes, skeletal myocytes or breast myoepithelial cells, e.g., *SYNM* (Figure 7 F.v) and *TPM1* (Figure 7 F.vi).

## DISCUSSION

Here we present a tissue-centric, cell type gene enrichment atlas, generated from the analysis of hundreds of biological replicates. Although it is frequently stated that cell-type gene expression profiles cannot be extracted from bulk RNAseq, e.g., (Li and Li, 2018; Mou et al., 2020), here we have identified cell type enriched or co-enriched genes, and charted temporal transcriptome changes underlying cell type differentiation. We made comparisons between cell type enrichment signatures across tissues, without the requirement for normalisation or batch effect adjustments, a significant issue when handling scRNAseq datasets, for which currently no universal solution (Chu et al., 2022; Mandelboum et al., 2019). Our analysis included cell types that are difficult to extract from tissue, e.g., adipocytes, and those that are sensitive to processing, e.g., kidney podocytes; issues that can hinder analysis (Denisenko et al., 2020; Rondini and Granneman, 2020), but are circumvented here as cell removal from tissue was not required. We identify lowly expressed transcripts as cell type enriched, many of which can be detected only in a small minority of cells annotated as a given type by scRNAseq, possibly due to limited read depth and high number of drop-out events (Hicks et al., 2018). Transcript level alone is not sufficient to predict protein levels (Liu et al., 2016) and so potential function of proteins encoded by such genes may have been overlooked.

Our study is the only cell type gene enrichment atlas generated independently of scRNAseq. Comparison of scRNAseq datasets generated from analysis of the same tissue type can reveal surprisingly low agreement between studies (Jiang et al., 2022; Squair et al., 2021), possibly due to the low number of samples typically analysed, and the associated lack of biological varience. For top cell type enriched genes in adipose tissue, agreement between data generated using our analysis method and several scRNAseq studies was equivalent or greater than between the scRNAseq studies themselves (Norreen-Thorsen et al., 2022). This could reflect the large sample set analysed and the associated biological variance represented. Our method also has scope for well-powered comparisons of cell type enrichment profiles between healthy and disease states, sexes (Norreen-Thorsen et al., 2022), ages, developmental stages, or metabolic states, using existing RNAseq resources for which phenotypic data is available, such as GTEx (Consortium, 2020) or TCGA (https://www.cancer.gov/tcga).

Various deconvolution algorithms have been developed to determine proportions of constituent cell types in bulk RNAseq, e.g., CIBERSORT and others (Glastonbury et al., 2019; Jew et al., 2020; Newman et al., 2015). Such analyses typically depend on input expression matrices of cell type reference genes, generated from transcriptome analysis of isolated cells or cell types. The accuracy of input matrices can be affected by various factors, such as technical artifacts due to cell extraction and processing, the presence of contaminating cell types, and limited input data availability for some cell types. Cross checking input matrices against our dataset could optimise such analysis, by identifying the most likely highly enriched genes *in vivo*.

### Limitations of the study

There are limitations to our study. In some tissues, we do not profile specific cell subtypes, e.g., basal and suprabasal keratinocytes in the skin, which were handled as one cell type in our analysis. In such cases, we failed to identify genes that fulfilled the criteria for use as input *Ref.T*.. In keeping with our observations, scRNAseq analysis of skin showed that genes considered to be basal keratinocyte markers e.g., *COL17A1* and *KRT5*, were indeed most highly expressed in this cell type, but were also co-enriched within the tissue in suprabasal keratinocytes (Karlsson et al., 2021). Thus, such cell subtype definitions are likely primarily governed by variation in absolute mRNA expression levels, rather that the presence or absence of a large number of uniquely enriched genes.

As our analysis end point is a gene enrichment score, we do not provide information on absolute mRNA expression profiles on a cell type basis, such as that generated by scRNAseq analysis.

As the prediction of cell type enriched genes is dependent on known input *Ref.T*., we cannot identify novel cell (sub)types for which *Ref.T*. have not yet been described.

We analysed samples from a total of 933 individuals from the GTEx portal (Consortium, 2020), with diverse health status, whose ages skewed older (ages 20-29: 8.5%, 30-39: 8.1%, 40-49: 15.6%, 50-59: 31.9%, 60-69: 32.4%, 70-79: 3.4%). Thus, the input dataset represents a limited age demographic, and a health status that may not represent the general population.

Expression of certain genes are strongly modified by environmental (e.g., eating, exercise, inflammation etc.), or genetic factors (Gibson, 2008). Such genes may therefore lack correlation with the constitutively expressed *Ref.T*. selected to represent the cell type in which they are predominantly transcribed, and thus could be considered a type of false negative in our analysis. One such example is *SELE*, an endothelial cell specific gene that is highly upregulated during inflammation and expressed at very low levels, if at all, in resting state (Vestweber and Blanks, 1999). *SELE*, despite its highly endothelial cell restricted expression profile, is not classified as EC enriched in our analysis, due to the variable nature of its expression.

We used relatively high thresholds for classification of cell type enriched genes. It is likely that some cell type enriched genes may be false negatives in our analysis, as they fall just below the thresholds required for classification as such. For example, the gene *KANK3* is classified as endothelial cell enriched in 9 tissue types, in the remaining 6 the highest enrichment score is also in endothelial cells, vs. all other types profiled, although it did not reach classification threshold. Thus, our classifications are intended only as a guide, and the reader should consider the data on a transcript-by-transcript basis.

All data generated in this study is available on the Human Protein Atlas in the ’Tissue Cell Type’ section (www.proteinatlas.org/humanproteome/tissue+cell+type), and can be used alongside data generated from scRNAseq in the ’Single Cell Type Section’ (Karlsson et al., 2021), and the antibody based tissue protein profiling in the ’Tissue Section’ (Uhlen et al., 2015).

## METHODS AND RESOURCES

### LEAD CONTACT AND MATERIALS AVAILABILITY

Further information and requests for resources and reagents should be directed to and will be fulfilled by the Lead Contact: Dr. Lynn Marie Butler.

This study did not generate new unique reagents.

### EXPERIMENTAL MODEL AND SUBJECT DETAILS

Bulk RNAseq data analysed in this study was obtained from the Genotype-Tissue Expression (GTEx) Project (gtexportal.org) V8 (Consortium, 2015) on 2021/04/26 (dbGaP Accession phs000424.v8.p2). Protein coding genes were categorised according to Biotype definitions in ENSEMBL release 102 (Yates et al., 2020) inclusive of those defined as “protein_coding”, “IG_C_gene”, “IG_D_gene”, “IG_J_gene”, “IG_V_gene”, “TR_C_gene”,”TR_D_gene”, ”TR_J_gene” and “TR_V_gene”. All other categorisations were classified as “non-protein coding” and were excluded from the analysis. Human tissue protein profiling was performed in house as part of the Human Protein Atlas project (Ponten et al., 2008; Uhlen et al., 2015; Uhlen et al., 2017) (www.proteinatlas.org). Normal tissue samples were obtained from the Department of Pathology, Uppsala University Hospital, Uppsala, Sweden, as part of the Uppsala Biobank. Samples were handled in accordance with Swedish laws and regulations, with approval from the Uppsala Ethical Review Board (Uhlen et al., 2015).

## METHOD DETAILS

### Sample inclusion

All samples in each GTEx tissue type dataset were included in the analyses, with the exception of: (i) ***Skin-not Sun Exposed (suprapubic)***: *Ref.T*. selected to represent hair root cells were absent or very lowly expressed in a large number of samples, presumably due to the lack of such structures in the selected region of tissue analysed. Thus, only samples with mean TPM >0.1 for hair follicle expressed transcripts trichohyalin (*TCHH*), keratins 25 (*KRT25*) and 71 (*KRT71*) were included for in the analysis (n=177). (ii) ***Breast – Mammary Tissue***: The GTEx breast dataset contains samples from both male and female donors, we analysed those from only from females. In both cases, sample IDs included in the analysis can be found in Table S2, Tab ’Sample IDs’.

## QUANTIFICATION AND STATISTICAL ANALYSIS

### Reference transcript-based correlation analysis

This method was based on that we previously developed (Butler et al., 2016; Dusart et al., 2019; Norreen-Thorsen et al., 2022). Pairwise Spearman correlation coefficients between reference transcripts (*Ref.T*.), selected as proxy markers for each cell type (see Table S1, Tab 1, Table A-O), and all other transcripts were calculated in R using the *corr.test* function from the *psych* package (v 1.8.4). False Discovery Rate (FDR) adjusted p-values (using Bonferroni correction) <0.0001 were considered significant. Genes were predicted to be cell type enriched if they fulfilled the criteria as described in the results section. In cases where a given cell type was represented by more than one *Ref.T* panel, or they could be considered related sub-cell types, the minimum differential score required vs. other *Ref.T*. panels was calculated excluding each the other (i.e., genes that correlated highly with both *Ref.T*. panels representing the same (sub)cell type were *not* excluded from classification as cell type enriched, but included in both – see Table S1, Tab 2).

### Weighted correlation network (WGCNA) analysis

The R package WGCNA (Langfelder and Horvath, 2008) was used to perform co-expression network analysis for gene clustering, on log2 expression TPM values. The analysis was performed according to recommended conditions in the WGCNA manual. Non-protein coding transcripts and transcripts with too many missing values were excluded using the goodSamplesGenes() function.

### Gene Ontology

The Gene Ontology Consortium (Ashburner et al., 2000) and PANTHER classification resource (Mi et al., 2013; Mi et al., 2016) were used to identify over represented terms in gene lists from the GO ontology (release date 2022-07-01) or reactome (release date 2021-10-01) databases. Plots of GO terms were created using the Clusterprofiler package in R (Wu et al., 2021) or REVIGO (Supek et al., 2011), as specified.

### Additional datasets and analysis

Single cell RNAseq data from Tabula Sapiens (Tabula Sapiens et al., 2022) was downloaded and UMAP plots created using the Seurat package in R (Hao et al., 2021). Human testis scRNAseq data was sourced from the human testis atlas (Guo et al., 2018). Tissue enriched genes were downloaded from the Human Protein Atlas (HPA) tissue atlas (Uhlen et al., 2015) or GTEx database (Consortium, 2015), as collated in the Harminozome database (Rouillard et al., 2016).

### Tissue Profiling: Human tissue sections

Immunohistochemistry (IHC) stained tissue sections were stained, as previously described (Ponten et al., 2008; Uhlen et al., 2015). Briefly, formalin fixed and paraffin embedded tissue samples were sectioned, de-paraffinised in xylene, hydrated in graded alcohols and blocked for endogenous peroxidase in 0.3% hydrogen peroxide diluted in 95% ethanol. For antigen retrieval, a Decloaking chamber® (Biocare Medical, CA) was used. Slides were boiled in Citrate buffer®, pH6 (Lab Vision, CA). Primary antibodies and a dextran polymer visualization system (UltraVision LP HRP polymer®, Lab Vision) were incubated for 30 min each at room temperature and slides were developed for 10 minutes using Diaminobenzidine (Lab Vision) as the chromogen. Slides were counterstained in Mayers hematoxylin (Histolab) and scanned using Scanscope XT (Aperio). Primary antibodies used for IHC staining are listed in Table S7.

### Other visualisation and analysis tools

Graphs and plots were made using Graphpad prism or the ggplot2 package in R (Wickham, 2016), unless otherwise specified. Circular plots were constructed using the R package *circlize* (Gu et al., 2014) and pubmed data was extracted using the easyPubMed package in R (https://CRAN.R-project.org/package=easyPubMed). Some figure illustrations were created using BioRender.com.

## DATA AVAILABILITY

Data for all protein coding genes and antibody-based protein profiling is provided on the Human Protein Atlas (Tissue Cell Type section) (www.proteinatlas.org/humanproteome/tissue+cell+type). This article also includes all individual tissue datasets generated (Table S2) and cell type enrichment categorisations (Table S1, Tab 4).

## AUTHOR CONTRIBUTIONS

Conceptualisation: LMB. Methodology: PD, LMB. Formal analysis: PD, SO, ES, MNT, MJI, LMB. Investigation: PD, SO, ES, MNT, MJI, FP, CL, LMB. Resources: FP, CL, JO, MU, LMB. Writing – Original Draft: PD, LMB. Writing – Review & Editing: All, Visualisation: PD, LMB. Supervision: LMB, PD. Funding Acquisition: JO, FP, CL, MU, LMB.

## ACKNOWLEDGEMENTS

This work was supported by funding granted to LMB from Hjärt Lungfonden (20170759, 20170537) and the Swedish Research Council (2019-01493). Main funding for the Human Protein Atlas was provided from the Knut and Alice Wallenberg Foundation (WCPR) and the Erling Persson Foundation (KCAP). We acknowledge the staff of the Human Protein Atlas program and the Science for Life Laboratory (SciLifeLab) for their valuable contributions. **Data usage:** We used data from the Genotype-Tissue Expression (GTEx) Project (gtexportal.org) (Consortium, 2015), supported by the Office of the Director of the National Institutes of Health, and by NCI, NHGRI, NHLBI, NIDA, NIMH, and NINDS.

## SUPPLEMENTAL TABLE LEGENDS

**Table S1. Reference transcript selection and analysis summary**.

(Tab 1): Correlation coefficient values were calculated between selected Ref.T. for constituent cell types in 15 tissues, presented as correlation matrices for each tissue. (Tab 2): Cell subtypes represented by more than one Ref.T panel within a single tissue, that were not excluded when calculating duel enrichment, as well as equivalent cell type names in the Tabula Sapiens and HPA databases. (Tab 3): Correlation and differential thresholds used within each cell type for classifying enriched genes. As well as the total number and percentage of V high, high and moderately enriched genes within each cell type. (Tab 4): Cell type enrichment predictions for protein coding gene expression in all 15 analysed tissues.

**Table S2. Sample IDs and Cell type correlation values**.

(Tab: Sample IDs): Sample IDs for the GTEx samples used to calculate RNA correlation values within each tissue. (Other tabs): Mean TPM, mean correlation values and differential values for all protein coding genes within each cell type and each tissue.

**Table S3. GO terms in Alpha and Beta cells of pancreas**

(Tab 1): Enriched GO Biological Process, Reactome, and Cellular Component classes for the list of 131 genes co-enriched in both Alpha and Beta cells of the pancreas. (Tab 2): Genes with predicted enrichment within both Alpha and Beta cells of the pancreas with functions linked to synapses.

**Table S4. Values and GO analysis of germ cell enriched genes**

(Tab 1): Cell type correlation and differential values of genes enriched within germ cell types within the testis. (Tab 2): Enriched GO Biological Process and reactome classes for the list of all germ cell enriched genes. (Tab 3): Enriched GO Biological Process classes for genes within each germ cell subtype separately.

**Table S5. GO analysis of genes enriched in multiple cell types**

(Tab 1): Table A: List of genes enriched in 3 or more of: adipocytes, sebaceous gland cells, hepatocytes and proximal tubular cells. Table B: Enriched GO Biological Process classes for genes listed in Table A. (Tab 2): Table A: List of genes enriched in both Respiratory Ciliated cells of the lung and either S3 or S4 cells (Early or Late Spermatids) of the testis. Table B: Enriched GO Biological Process classes for genes listed in Table A. (Tab 3): Table A: List of genes enriched in either S3 or S4 cells (Early or Late Spermatids) of the testis, but NOT enriched in Respiratory Ciliated cells of the lung. Table B: Enriched GO Biological Process classes for genes listed in Table A.

**Table S6. Core cell type enriched genes**

Lists of genes predicted to be cell type enriched in at least half the tissues profiled for each core cell type.

**Table S7. Primary antibodies**

List of IDs for all primary antibodies used to stain all immunohistochemistry images used in this study.

