## Supplemental Figures for "A tissue centric atlas of cell type transcriptome enrichment signatures"

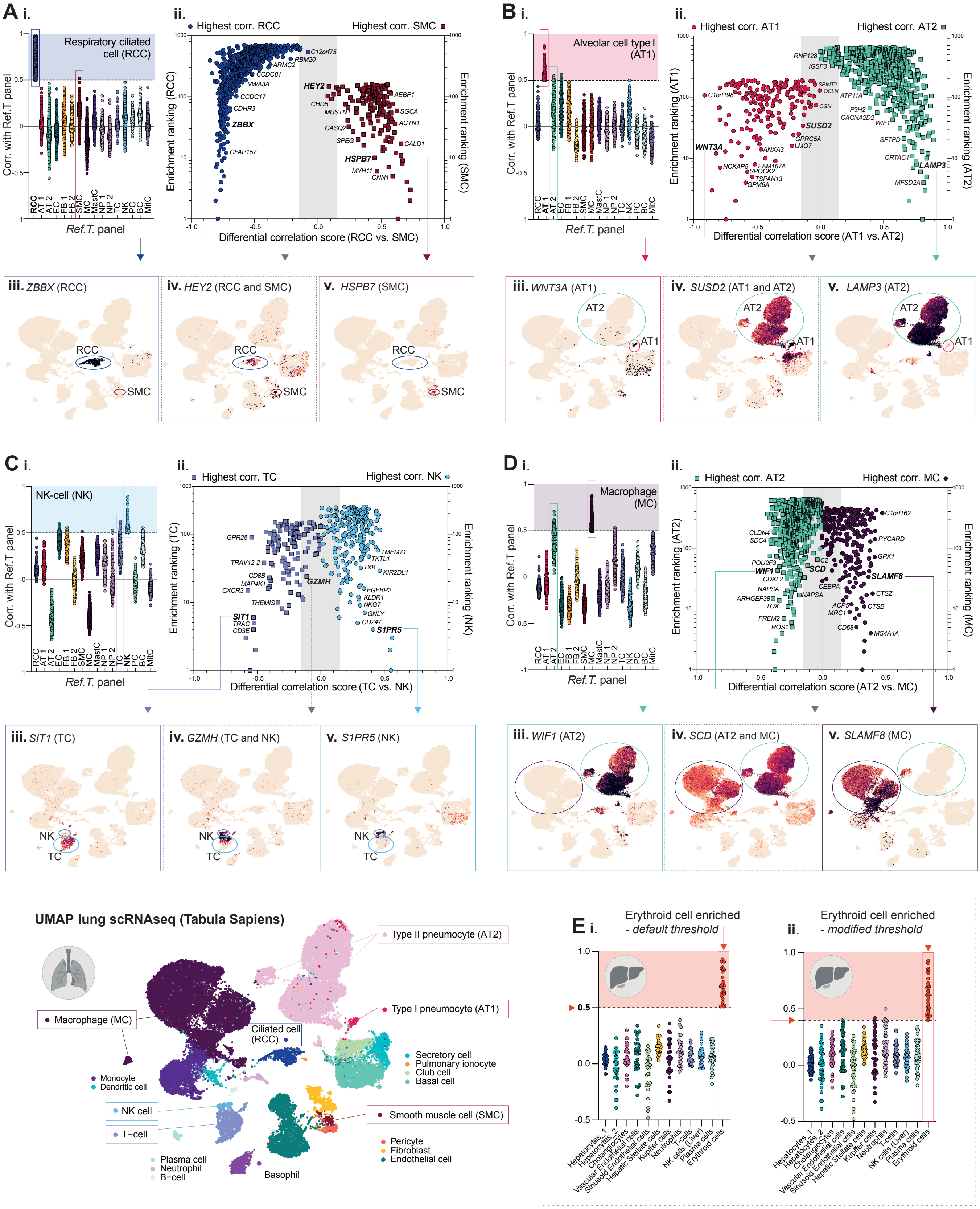

A i.
Respiratory ciliated cell (RCC)
Corr. with Ref.T panel
Ref.T. panel
ii.
Highest corr. RCC
Highest corr. SMC
Enrichment ranking (RCC)
Enrichment ranking (SMC)
Differential correlation score (RCC vs. SMC)
iii. ZBBX (RCC)
iv. HEY2 (RCC and SMC)
v. HSPB7 (SMC)
B i.
Alveolar cell type I (AT1)
ii.
Highest corr. AT1
Highest corr. AT2
Enrichment ranking (AT1)
Enrichment ranking (AT2)
Differential correlation score (AT1 vs. AT2)
iii. WNT3A (AT1)
iv. SUSD2 (AT1 and AT2)
v. LAMP3 (AT2)
C i.
NK-cell (NK)
ii.
Highest corr. TC
Highest corr. NK
Enrichment ranking (TC)
Enrichment ranking (NK)
Differential correlation score (TC vs. NK)
iii. SIT1 (TC)
iv. GZMH (TC and NK)
v. S1PR5 (NK)
D i.
Macrophage (MC)
ii.
Highest corr. AT2
Highest corr. MC
Enrichment ranking (AT2)
Enrichment ranking (MC)
Differential correlation score (AT2 vs. MC)
iii. WIF1 (AT2)
iv. SCD (AT2 and MC)
v. SLAMF8 (MC)
UMAP lung scRNAseq (Tabula Sapiens)
Type II pneumocyte (AT2)
Type I pneumocyte (AT1)
Macrophage (MC)
Ciliated cell (RCC)
Secretory cell
Pulmonary ionocyte
Club cell
Basal cell
Monocyte
Dendritic cell
NK cell
T-cell
Smooth muscle cell (SMC)
Pericyte
Fibroblast
Endothelial cell
Plasma cell
Neutrophil
B-cell
Basophil
E i.
Erythroid cell enriched - default threshold
ii.
Erythroid cell enriched - modified threshold
Hepatocytes_1
Hepatocytes_2
Cholangiocytes
Vascular Endothelial cells
Sinusoid Endothelial cells
Hepatic Stellate cells
Kupffer cells
Neutrophils
T-cells
NK cells (Liver)
Plasma cells
Erythroid cells

**Figure S1. Integrative co-expression analysis of unfractionated human tissue RNAseq can resolve constituent cell type enriched genes. Related to Figure 1.**

RNAseq datasets for human lung (n=578) were retrieved from GTEx V8 and correlation coefficients between selected cell type *Ref. T.* and all other sequenced transcripts generated. Correlation values vs. all other cell type *Ref.T.* panels for transcripts reaching the designated threshold with *Ref. T.* for (**A**) (i) respiratory ciliated cells (RCC) (**B**) (i) alveolar type I cells (AT1), (**C**) (i) natural killer cells (NK) or (**D**) (i) macrophages (MC). The '*differential correlation score'* and respective enrichment rankings for transcripts reaching the designated threshold with *Ref. T.* for (**A**) (ii) RCC or SMC, (**B**) (ii) AT1 or AT2, (**C**) (ii) NK or TC and (**D**) (ii) MC and AT2. scRNAseq data from analysis of human lung was sourced from Tabula Sapiens (Tabula Sapiens et al., 2022) and used to generate UMAP plots, showing the expression profiles of example genes we predicted as being enriched in (**A**) (iii) RCC only, (iv) RCC and SMC or (v) SMC only, (**B**) (iii) AT1 only, (iv) AT1 and AT2 or (v) AT2 only, (**C**) (iii) TC only, (iv) TC and NK or (v) NK only, or (**D**) (iii) AT2 only, (iv) AT2 and MC or (v) MC only. (**E**). RNAseq datasets for human liver (n=226) were retrieved from GTEx V8 and analysed as described for lung. Correlation values vs. all cell type *Ref.T.* panels for transcripts reaching the (i) designated or (ii) modified threshold for classification as erythroid cell enriched. EC; Endothelial cell, FB1/FB2; fibroblast, MC; macrophage, MastC; mast cell, NP1/NP2; neutrophil, TC; T-cell, NK; natural killer cell, PC; plasma cell, BC; B-cell.

### Figure S2

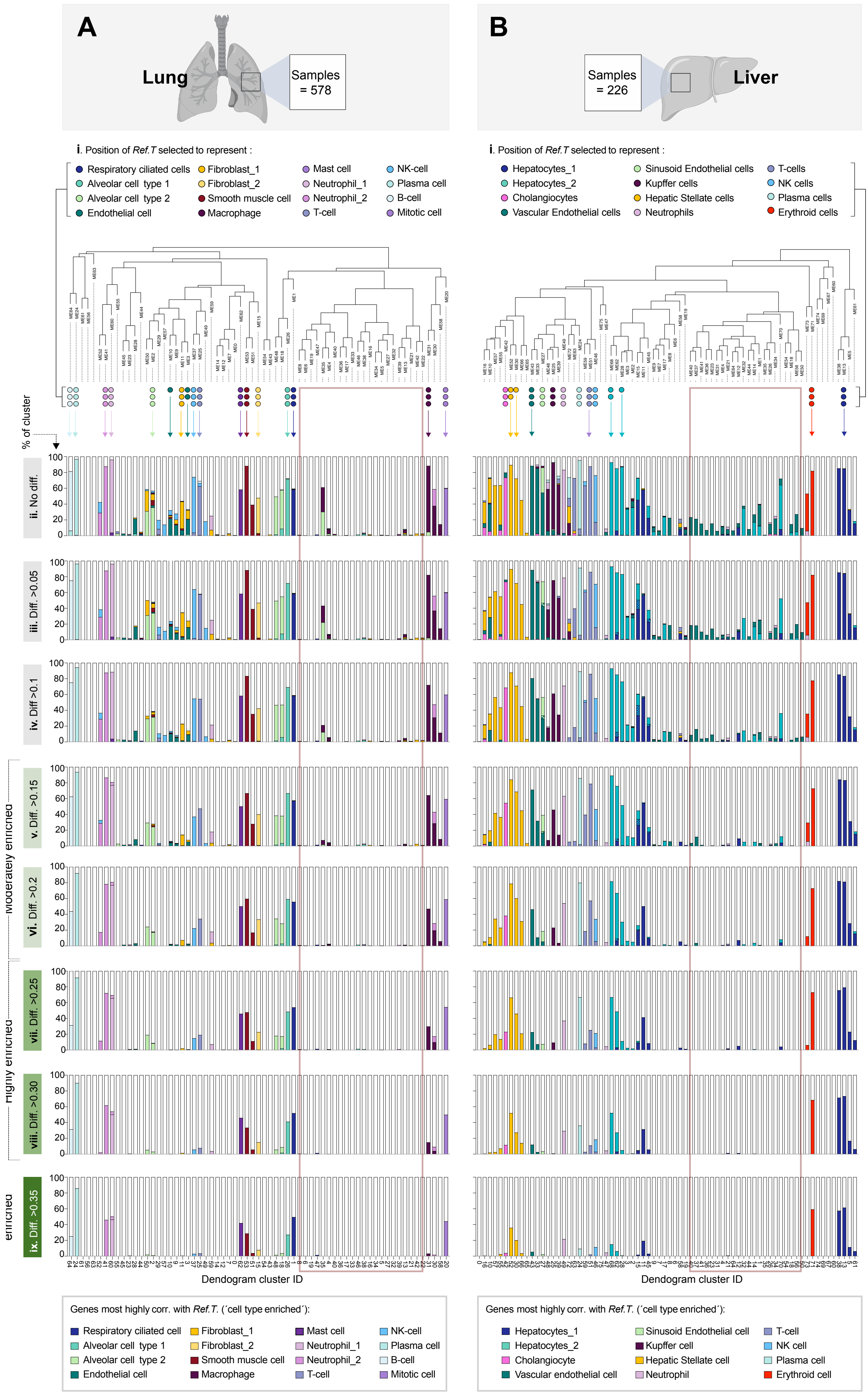
A
Lung
Samples = 578
B
Samples = 226
Liver
i. Position of Ref.T selected to represent :
Respiratory ciliated cells
Alveolar cell type 1
Alveolar cell type 2
Endothelial cell
Fibroblast_1
Fibroblast_2
Smooth muscle cell
Macrophage
Mast cell
Neutrophil_1
Neutrophil_2
T-cell
NK-cell
Plasma cell
B-cell
Mitotic cell
Hepatocytes_1
Hepatocytes_2
Cholangiocytes
Vascular Endothelial cells
Sinusoid Endothelial cells
Kupffer cells
Hepatic Stellate cells
Neutrophils
T-cells
NK cells
Plasma cells
Erythroid cells
% of cluster
ii. No diff.
iii. Diff. >0.05
iv. Diff >0.1
v. Diff. >0.15
vi. Diff. >0.2
vii. Diff. >0.25
viii. Diff. >0.30
ix. Diff. >0.35
Moderately enriched
Highly enriched
enriched
Dendogram cluster ID
Dendogram cluster ID
Genes most highly corr. with Ref.T. (´cell type enriched´):
Respiratory ciliated cell
Alveolar cell type 1
Alveolar cell type 2
Endothelial cell
Fibroblast_1
Fibroblast_2
Smooth muscle cell
Macrophage
Mast cell
Neutrophil_1
Neutrophil_2
T-cell
NK-cell
Plasma cell
B-cell
Mitotic cell
Genes most highly corr. with Ref.T. (´cell type enriched´):
Hepatocytes_1
Hepatocytes_2
Cholangiocyte
Vascular endothelial cell
Sinusoid Endothelial cell
Kupffer cell
Hepatic Stellate cell
Neutrophil
T-cell
NK cell
Plasma cell
Erythroid cell

**Figure S2. Unsupervised weighted network correlation analysis (WGNCA) is consistent with *Ref.T.* analysis. Related to Figure 1.** RNAseq data from human (**A**) lung (n=578 individuals) or (**B**) pancreas (n=328) was subject to weighted correlation network analysis (WGCNA). In the resultant dendrograms, the position of (i) *Ref.T*. selected to represent each cell type and (ii) the % of the cluster containing transcripts that had a correlation with any *Ref.T*. panel above the designated threshold, are indicated; colour representing the cell type classification (see bottom panel) (Table S1, Tab 5 for thresholds). Distribution of transcripts for each cell type classification when the highest correlation with any given *Ref.T*. panel was a minimum of (ii) 0, (iii) 0.05, (iv) 0.10, (v) 0.15 [moderately enriched], (vi) 0.20, (vii) 0.25 [highly enriched] or (viii) 0.30 or (ix) 0.35 [very highly enriched] greater that the next highest with a different *Ref.T.* panel (´differential correlation score´).

Figure S3

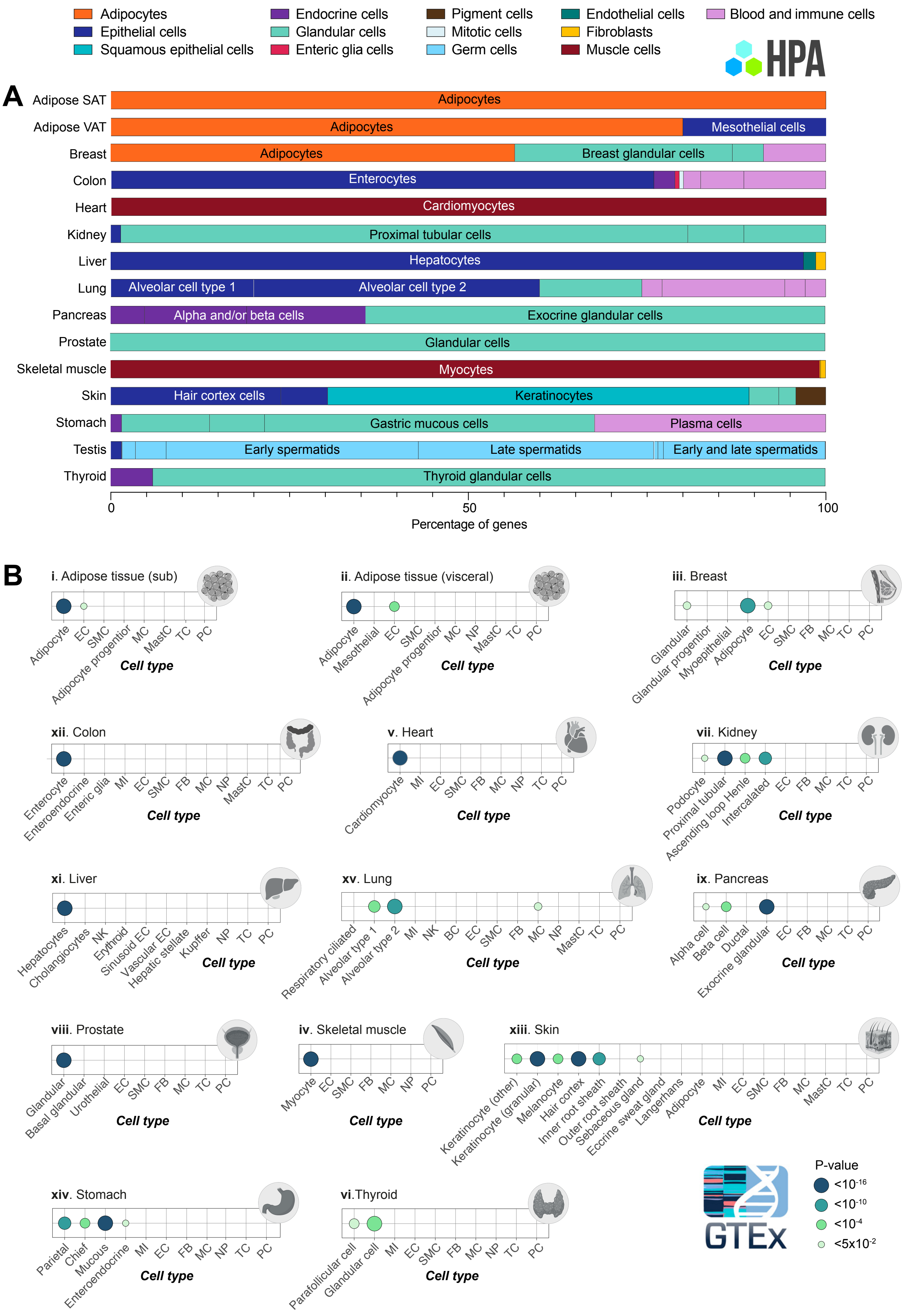

Adipocytes
Epithelial cells
Squamous epithelial cells
Endocrine cells
Glandular cells
Enteric glia cells
Pigment cells
Mitotic cells
Germ cells
Endothelial cells
Fibroblasts
Muscle cells
Blood and immune cells
HPA
A
Adipose SAT
Adipose VAT
Breast
Colon
Heart
Kidney
Liver
Lung
Pancreas
Prostate
Skeletal muscle
Skin
Stomach
Testis
Thyroid
Adipocytes
Mesothelial cells
Breast glandular cells
Enterocytes
Cardiomyocytes
Proximal tubular cells
Hepatocytes
Alveolar cell type 1
Alveolar cell type 2
Alpha and/or beta cells
Exocrine glandular cells
Glandular cells
Myocytes
Hair cortex cells
Keratinocytes
Gastric mucous cells
Plasma cells
Early spermatids
Late spermatids
Early and late spermatids
Thyroid glandular cells
0
50
100
Percentage of genes
B
i. Adipose tissue (sub)
ii. Adipose tissue (visceral)
iii. Breast
xii. Colon
v. Heart
vii. Kidney
xi. Liver
xv. Lung
ix. Pancreas
viii. Prostate
iv. Skeletal muscle
xiii. Skin
xiv. Stomach
vi.Thyroid
Cell type
P-value
<10-16
<10-10
<10-4
<5x10-2
GTEx

**Figure S3. Integrative co-expression analysis of unfractionated human tissue RNAseq can resolve tissue enriched genes into single cell type expression source. Related to Figure 2.** (**A**) Bar plot showing the fraction of predicted cell type enriched genes among the tissue, or tissue-group, enriched genes in Human Protein Atlas (HPA). Colour indicates cell type group. The cell type with the most shared enriched genes with tissues are labelled. (B) Bubble plots showing the significance (indicated by dot size and colour) of similarity between the top 300 tissue enriched genes in GTEx and the predicted cell type enrichment signatures Where overlap is not statistically significant (hypergeometric test, $P > 0.05$), the corresponding dot is removed. EC; endothelial cell, SMC; Smooth muscle cell, MC; macrophage, MastC; mast cell, TC; T-cell, PC; Plasma cell, NP; Neutrophil, MI; Mitotic cell, NK; Natural killer cell.

Figure S4

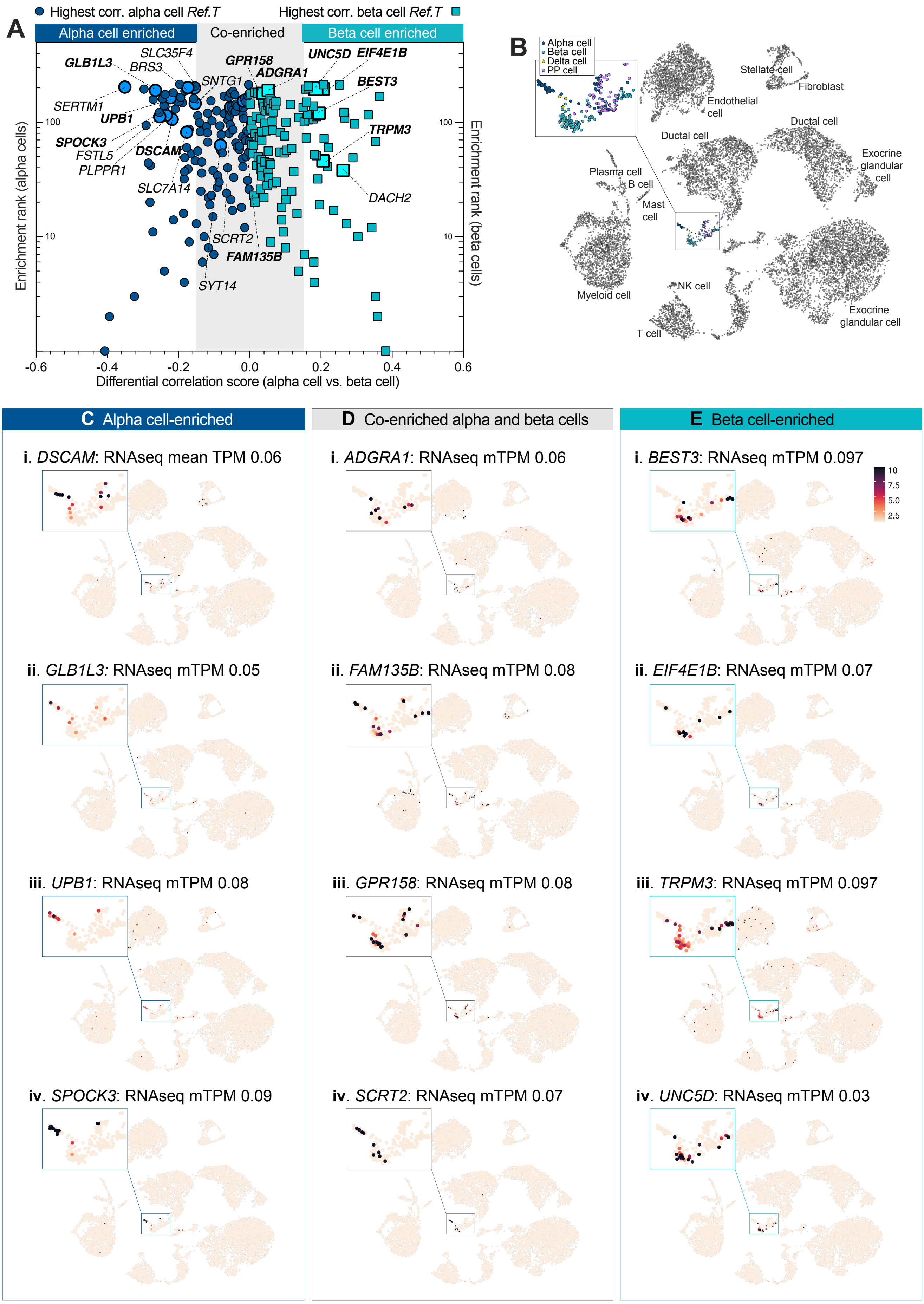
A
Highest corr. alpha cell Ref.T
Highest corr. beta cell Ref.T
Alpha cell enriched
Co-enriched
Beta cell enriched
GLB1L3
SLC35F4
BRS3
GPR158
ADGRA1
UNC5D
EIF4E1B
SNTG1
BEST3
SERTM1
UPB1
TRPM3
SPOCK3
DSCAM
FSTL5
PLPPR1
SLC7A14
DACH2
SCRT2
FAM135B
SYT14
Enrichment rank (alpha cells)
Enrichment rank (beta cells)
100
10
-0.6
-0.4
-0.2
0.0
0.2
0.4
0.6
Differential correlation score (alpha cell vs. beta cell)
B
Alpha cell
Beta cell
Delta cell
PP cell
Stellate cell
Fibroblast
Endothelial cell
Ductal cell
Ductal cell
Exocrine glandular cell
Plasma cell
B cell
Mast cell
Myeloid cell
NK cell
T cell
Exocrine glandular cell
C Alpha cell-enriched
i. DSCAM: RNAseq mean TPM 0.06
ii. GLB1L3: RNAseq mTPM 0.05
iii. UPB1: RNAseq mTPM 0.08
iv. SPOCK3: RNAseq mTPM 0.09
D Co-enriched alpha and beta cells
i. ADGRA1: RNAseq mTPM 0.06
ii. FAM135B: RNAseq mTPM 0.08
iii. GPR158: RNAseq mTPM 0.08
iv. SCRT2: RNAseq mTPM 0.07
E Beta cell-enriched
i. BEST3: RNAseq mTPM 0.097
10
7.5
5.0
2.5
ii. EIF4E1B: RNAseq mTPM 0.07
iii. TRPM3: RNAseq mTPM 0.097
iv. UNC5D: RNAseq mTPM 0.03

**Figure S4. Reference transcript-based identification of lowly expressed pancreatic alpha and beta cell type enriched and co-enriched genes. Related to Figure 3.** RNAseq datasets for human pancreas (n=328) were analysed to generate correlation coefficient values between all protein coding genes and *Ref T*. (**A**) For genes that correlated most highly with alpha (dark blue) or beta cell (turquoise) *Ref.T* (above >0.50), the '*differential correlation score'* (difference between mean corr. with alpha and beta cell *Ref.T.*) was plotted vs. 'enrichment ranking' (position in each respective list, highest corr. = rank 1). Shaded grey box highlights genes enriched in both cell types (co-enriched). Genes highlighted in bold correspond to those featured in the lower panels. scRNAseq data from analysis of human pancreas was sourced from Tabula Sapiens (Tabula Sapiens et al., 2022), and used to generate UMAP plots showing (**B**) scRNAseq cell type annotations, and the expression profiles of genes we predicted as being (**C**) alpha cell-enriched; (i) *DSCAM*, (ii) *GLB1L3*, (iii) *UPB1* and (iv) *SPOCK3*, (**D**) co-enriched in both alpha and beta cells; (i) *ADGRA1*, (ii) *FAM135B*, (iii) *GPR158* and (iv) *SCRT2*, or (**E**) beta cell-enriched; (i) *BEST3*, (ii) *EIF4E1B*, (iii) *TRPM3* and (iv) *UNC5D.*

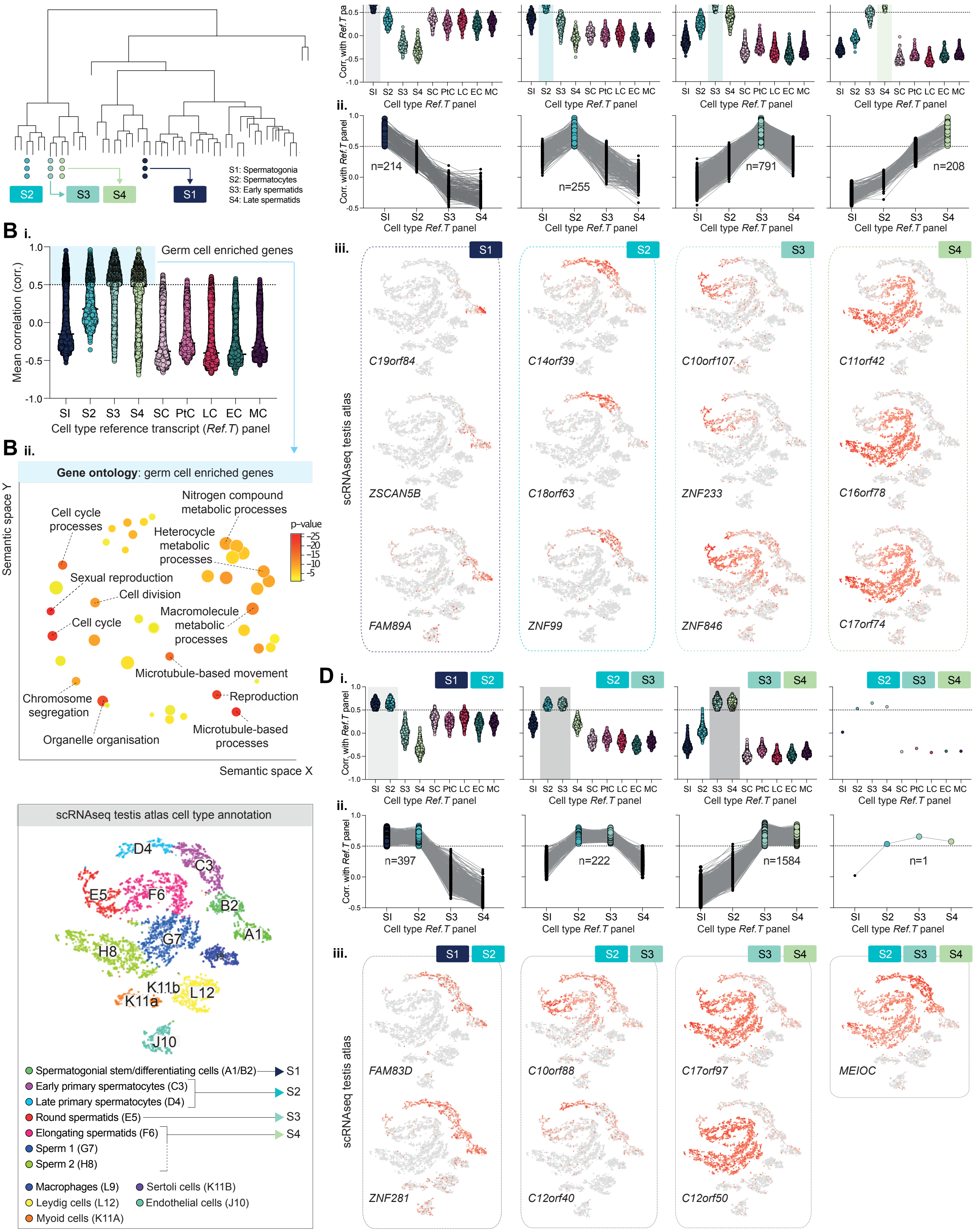

S1: Spermatogonia
S2: Spermatocytes
S3: Early spermatids
S4: Late spermatids
S2
S3
S4
S1
Corr. with Ref.T panel
Cell type Ref.T panel
SI S2 S3 S4 SC PtC LC EC MC
ii.
n=214
n=255
n=791
n=208
B i.
Germ cell enriched genes
Mean correlation (corr.)
Cell type reference transcript (Ref.T) panel
iii.
scRNAseq testis atlas
C19orf84
ZSCAN5B
FAM89A
C14orf39
C18orf63
ZNF99
C10orf107
ZNF233
ZNF846
C11orf42
C16orf78
C17orf74
B ii.
Gene ontology: germ cell enriched genes
Semantic space Y
Semantic space X
p-value
Cell cycle processes
Nitrogen compound metabolic processes
Heterocycle metabolic processes
Sexual reproduction
Cell division
Macromolecule metabolic processes
Cell cycle
Microtubule-based movement
Chromosome segregation
Reproduction
Microtubule-based processes
Organelle organisation
D i.
n=397
n=222
n=1584
n=1
FAM83D
ZNF281
C10orf88
C12orf40
C17orf97
C12orf50
MEIOC
scRNAseq testis atlas cell type annotation
D4
C3
E5
F6
B2
G7
A1
H8
K11b
L12
K11a
J10
Spermatogonial stem/differentiating cells (A1/B2)
Early primary spermatocytes (C3)
Late primary spermatocytes (D4)
Round spermatids (E5)
Elongating spermatids (F6)
Sperm 1 (G7)
Sperm 2 (H8)
Macrophages (L9)
Sertoli cells (K11B)
Leydig cells (L12)
Endothelial cells (J10)
Myoid cells (K11A)

**Figure S5. Analysis of pseudo temporal changes during spermatogenesis reveals stage-specific and common stage-shared gene enrichment signatures. Related to Figure 4.** (**A**) weighted network correlation analysis of human testis RNAseq data (n=361) annotated to show position of genes in *Ref.T.* panels (each indicated with single circle) selected to represent cell types at the different stages of spermatogenesis: S1 (spermatogonia), S2 (spermatocytes), S3 and S4 (early and late spermatids, respectively). (**B**) For genes with predicted cell-type enrichment in S1, S2, S3 or S4 (i) mean correlation coefficients with *Ref.T.* for S1, S2, S3, S4 and sertoli cells (SC), Leydig cells (LC), peritubular cells (PtC), endothelial cells (EC) or macrophages (MC) and (ii) over-represented gene ontology terms, summarised and visualised using REVIGO. For all genes predicted to be: (**C**) highly cell type enriched at one stage of spermatogenesis or (**D**) co-enriched at two or more stages of spermatogenesis (category indicated in top left of each plot): (i) mean correlation coefficients with *Ref.T.* for S1, S2, S3, S4, SC, LC, PtC, EC or MC, (ii) mean correlation coefficients with *Ref.T.* for S1, S2, S3, S4 with linkage lines connecting each individual gene (iii) expression profiles in Human Testis Atlas scRNAseq data (Guo et al., 2018) for selected lesser known genes appearing in each respective category. UMAP from the Human Testis Atlas shows original cell type annotations (bottom left), with arrows to indicating the broad equivalence classifications in our analysis.

### Figure S6

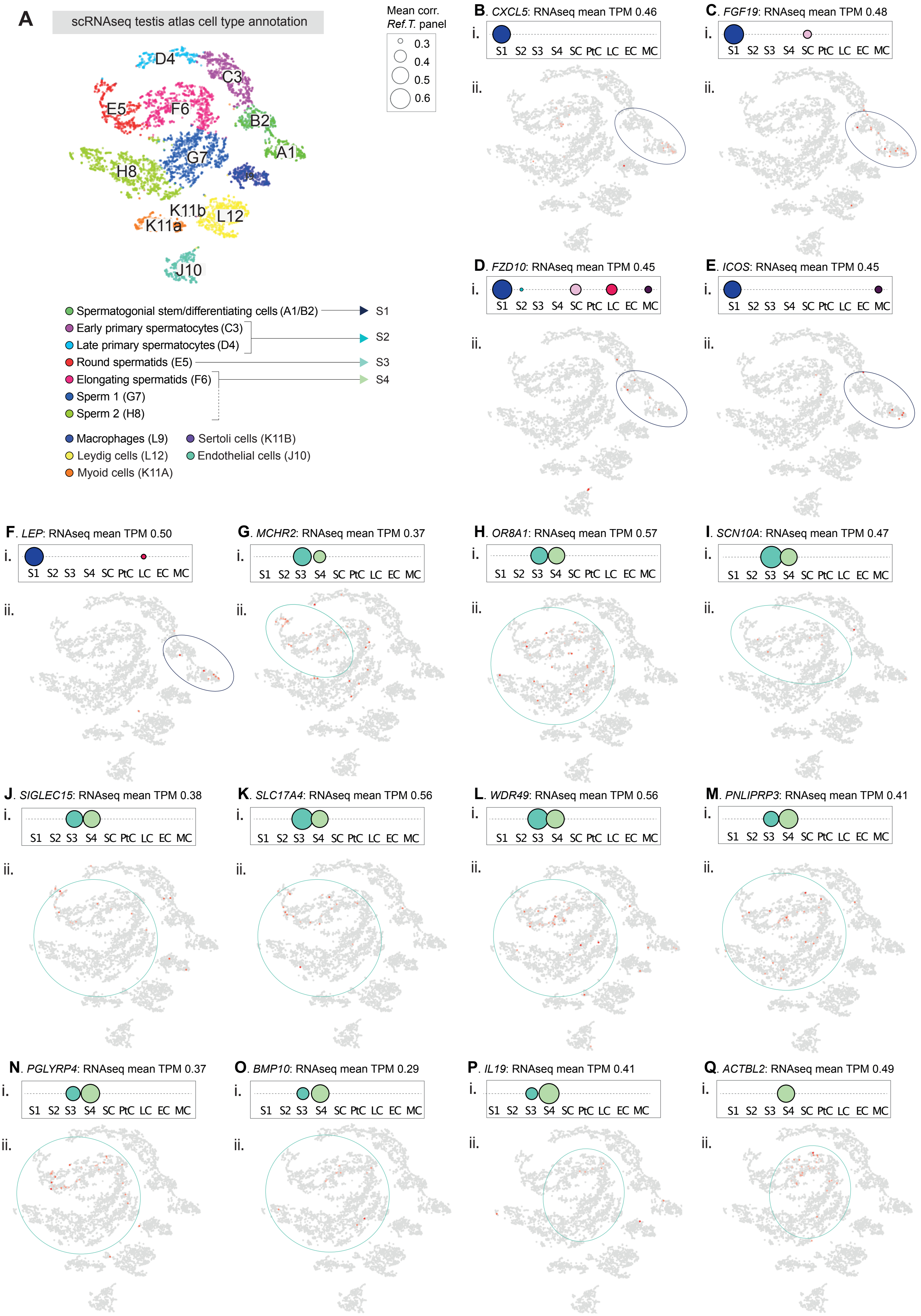

A
scRNAseq testis atlas cell type annotation
D4
C3
E5
F6
B2
A1
G7
H8
K11b
L12
K11a
J10
Spermatogonial stem/differentiating cells (A1/B2)
Early primary spermatocytes (C3)
Late primary spermatocytes (D4)
Round spermatids (E5)
Elongating spermatids (F6)
Sperm 1 (G7)
Sperm 2 (H8)
Macrophages (L9)
Leydig cells (L12)
Myoid cells (K11A)
Sertoli cells (K11B)
Endothelial cells (J10)
S1
S2
S3
S4
Mean corr.
Ref.T. panel
0.3
0.4
0.5
0.6
B. CXCL5: RNAseq mean TPM 0.46
C. FGF19: RNAseq mean TPM 0.48
D. FZD10: RNAseq mean TPM 0.45
E. ICOS: RNAseq mean TPM 0.45
F. LEP: RNAseq mean TPM 0.50
G. MCHR2: RNAseq mean TPM 0.37
H. OR8A1: RNAseq mean TPM 0.57
I. SCN10A: RNAseq mean TPM 0.47
J. SIGLEC15: RNAseq mean TPM 0.38
K. SLC17A4: RNAseq mean TPM 0.56
L. WDR49: RNAseq mean TPM 0.56
M. PNLIPRP3: RNAseq mean TPM 0.41
N. PGLYRP4: RNAseq mean TPM 0.37
O. BMP10: RNAseq mean TPM 0.29
P. IL19: RNAseq mean TPM 0.41
Q. ACTBL2: RNAseq mean TPM 0.49
i.
ii.
S1 S2 S3 S4 SC PtC LC EC MC

**Figure S6. Reference transcript-based identification of lowly expressed germ cell enriched genes in the human testis. Related to Figure 4.** (**A**) UMAP and cell type annotations as defined in the scRNAseq Human Testis Atlas (Guo et al., 2018), with arrows to indicate the broad equivalence classifications in our analysis. (i) Enrichment scores in all cell types profiled for genes predicted to be (**B-F**) S1 enriched, (**H-N**) S3 and S4 enriched or (**O-Q**) S4 enriched, with (ii) corresponding UMAP expression plots from the scRNAseq Human Testis Atlas (Guo et al., 2018).

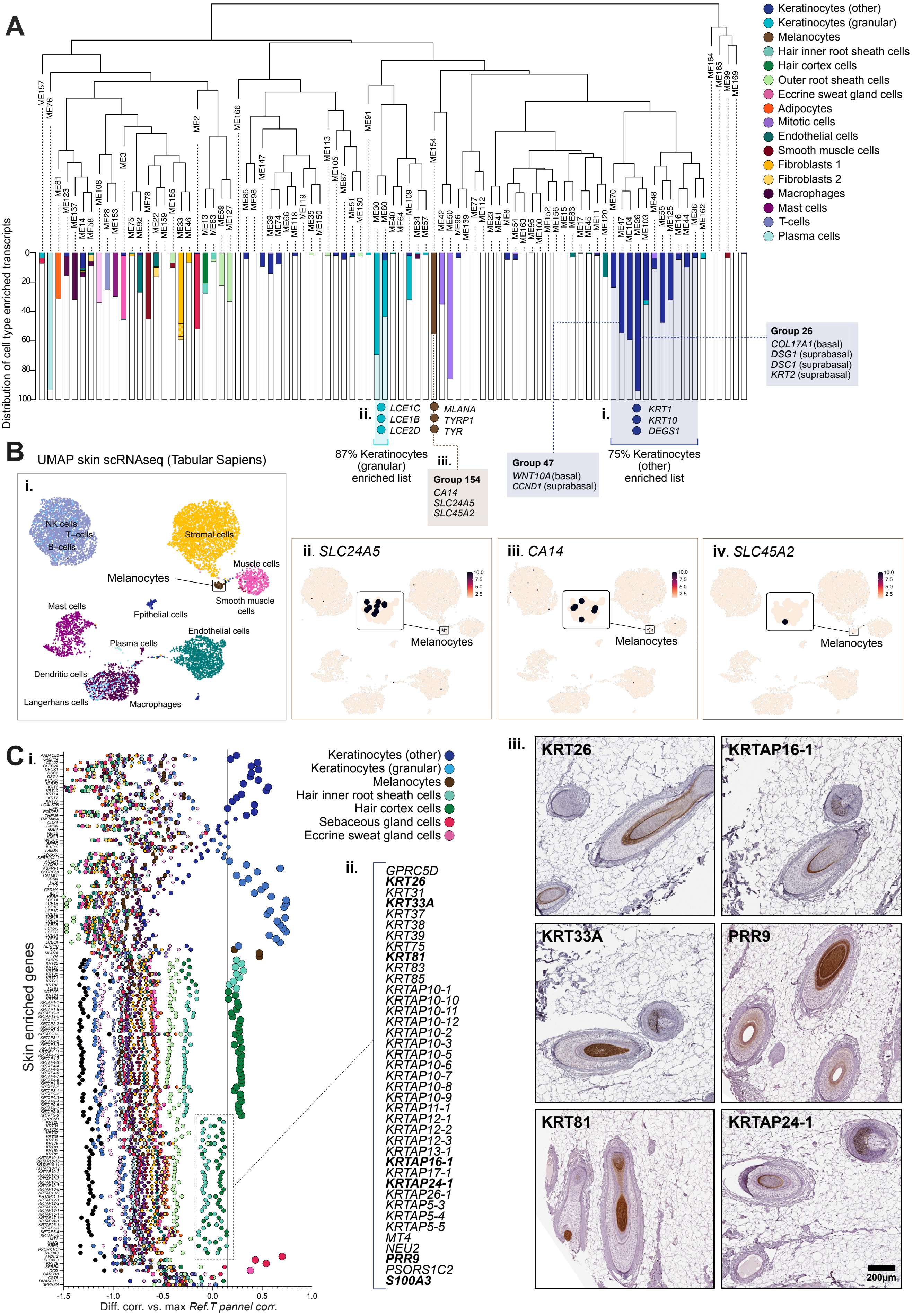
A
Keratinocytes (other)
Keratinocytes (granular)
Melanocytes
Hair inner root sheath cells
Hair cortex cells
Outer root sheath cells
Eccrine sweat gland cells
Adipocytes
Mitotic cells
Endothelial cells
Smooth muscle cells
Fibroblasts 1
Fibroblasts 2
Macrophages
Mast cells
T-cells
Plasma cells
Distribution of cell type enriched transcripts
Group 26
COL17A1 (basal)
DSG1 (suprabasal)
DSC1 (suprabasal)
KRT2 (suprabasal)
ii. LCE1C LCE1B LCE2D
MLANA TYRP1 TYR
i. KRT1 KRT10 DEGS1
87% Keratinocytes (granular) enriched list
iii.
Group 154
CA14
SLC24A5
SLC45A2
Group 47
WNT10A (basal)
CCND1 (suprabasal)
75% Keratinocytes (other) enriched list
B
UMAP skin scRNAseq (Tabular Sapiens)
i.
NK cells
T-cells
B-cells
Stromal cells
Muscle cells
Melanocytes
Smooth muscle cells
Mast cells
Epithelial cells
Endothelial cells
Plasma cells
Dendritic cells
Langerhans cells
Macrophages
ii. SLC24A5
iii. CA14
iv. SLC45A2
Melanocytes
C i.
Keratinocytes (other)
Keratinocytes (granular)
Melanocytes
Hair inner root sheath cells
Hair cortex cells
Sebaceous gland cells
Eccrine sweat gland cells
Skin enriched genes
Diff. corr. vs. max Ref.T pannel corr.
ii.
GPRC5D
KRT26
KRT31
KRT33A
KRT37
KRT38
KRT39
KRT75
KRT81
KRT83
KRT85
KRTAP10-1
KRTAP10-10
KRTAP10-11
KRTAP10-12
KRTAP10-2
KRTAP10-3
KRTAP10-5
KRTAP10-6
KRTAP10-7
KRTAP10-8
KRTAP10-9
KRTAP11-1
KRTAP12-1
KRTAP12-2
KRTAP12-3
KRTAP13-1
KRTAP16-1
KRTAP17-1
KRTAP24-1
KRTAP26-1
KRTAP5-3
KRTAP5-4
KRTAP5-5
MT4
NEU2
PRR9
PSORS1C2
S100A3
iii.
KRT26
KRTAP16-1
KRT33A
PRR9
KRT81
KRTAP24-1
200µm

**Figure S7. Constituent cells of the skin hair root are the primary source of skin tissue enriched genes. Related to Figure 5.** (**A**) Weighted network correlation analysis (WGNCA) of human skin samples (n=210) with coloured coded bars showing distribution of genes predicted to be cell type enriched. Position of *Ref.T.* and example cell-type enriched genes are highlighted for: (i) supra-basal keratinocytes, (ii) granular keratinocytes and (iii) melanocytes. (**B**) scRNAseq data and cell type definitions were sourced for human skin from Tabula Sapiens (Tabula Sapiens et al., 2022) and used to generate UMPA plots showing: (i) cell type annotations or expression profiles for genes we predicted to be melanocyte enriched (ii) *SLC24A5*, (iii) *CA14* and (iv) *SLC45A2*. (**C**) Skin enriched genes (vs. other tissue types) were identified and (i) corresponding cell type enrichment profiles in skin plotted, a panel of which (ii) did not reach the threshold for classification as enriched in a single cell type but had highest enrichment scores in one or more hair cell types. (iii) Expression of proteins encoded by selected examples were profiled in human skin tissue containing hair roots.

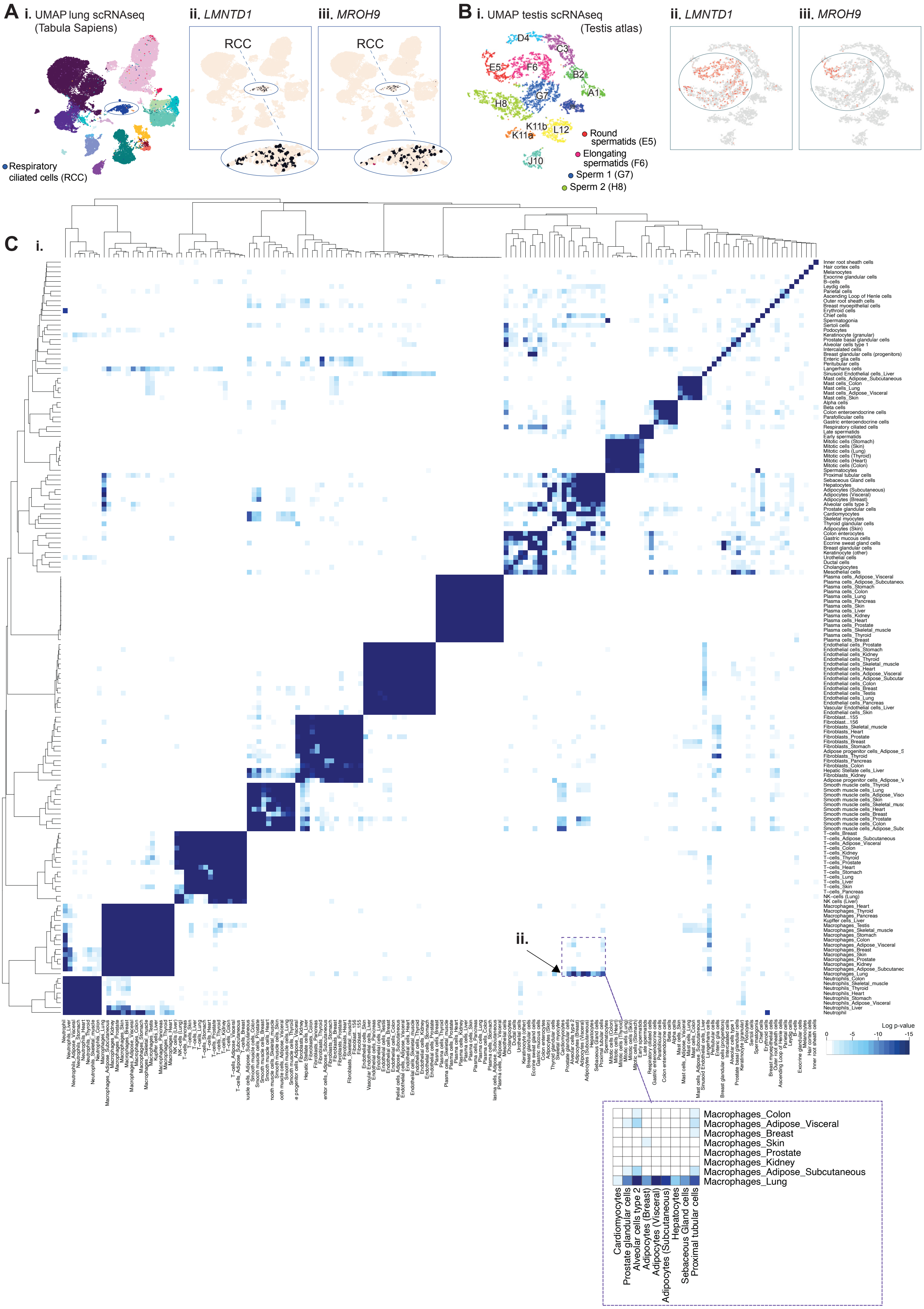
A i. UMAP lung scRNAseq (Tabula Sapiens)
Respiratory ciliated cells (RCC)
ii. LMNTD1
RCC
iii. MROH9
RCC
B i. UMAP testis scRNAseq (Testis atlas)
D4
C3
E5
F6
B2
G7
A1
H8
K11b
K11a
L12
J10
Round spermatids (E5)
Elongating spermatids (F6)
Sperm 1 (G7)
Sperm 2 (H8)
ii. LMNTD1
iii. MROH9
C i.
ii.
Log p-value
0
-5
-10
-15
Macrophages_Colon
Macrophages_Adipose_Visceral
Macrophages_Breast
Macrophages_Skin
Macrophages_Prostate
Macrophages_Kidney
Macrophages_Adipose_Subcutaneous
Macrophages_Lung
Cardiomyocytes
Prostate glandular cells
Alveolar cells type 2
Adipocytes (Breast)
Adipocytes (Visceral)
Adipocytes (Subcutaneous)
Hepatocytes
Sebaceous Gland cells
Proximal tubular cells

**Figure S8. Cell type enriched signature comparisons. Related to Figure 6 and 7.** scRNAseq data was sourced for human (**A**) lung from Tabula Sapiens (Tabula Sapiens et al., 2022) or (**B**) testis from the Human Testis Atlas (Guo et al., 2018), and used to generate UMAP plots to show (i) cell type annotation as according to the original studies, or expression profiles of (ii) *LMNTD1* or (iii) *MROH9*. (**C**) Heatmap showing significance p-values for similarity scores, calculated using a hypergeometric test, between: (i) all predicted cell type enriched genes, and (ii) lung macrophages vs. other non-macrophage cell types.
